# Unsaturated lipids as key control points for caveola formation and disassembly

**DOI:** 10.1101/2024.09.22.614381

**Authors:** Yeping Wu, Ye-Wheen Lim, Kerrie-Ann McMahon, Nick Martel, James Rae, Harriet P. Lo, Ya Gao, Vikas Tillu, Elin Larsson, Richard Lundmark, Daniel S. Levic, Michel Bagnat, Junxian Lim, David P. Fairlie, Albert Pol, Brett M. Collins, Nicholas Ariotti, Thomas E. Hall, Robert G. Parton

## Abstract

Caveolae are specialized plasma membrane domains with a unique lipid composition. Recent work has implicated lipid peroxidation in triggering caveola disassembly, releasing caveolar proteins to regulate the cellular response to oxidative stress. Here we investigated the role of specific lipid species in caveola formation and response to oxidative stress. Systematic screening of lipid enzymes identified ACSL4, a critical enzyme in the synthesis of polyunsaturated fatty acid-containing lipids, and enzymes involved in the biosynthesis of pro-ferroptotic ether lipids, as key regulators of caveola formation. Omega-6 fatty acids promoted efficient caveola formation, while their oxidation or displacement by omega-3 or monounsaturated fatty acids disrupted this process. Moreover, we found that omega-6 fatty acid-containing membrane lipids were required for oxidative stress-induced caveola disassembly. These findings unveil a new model for caveola formation, highlighting specific unsaturated lipids as essential caveolar components and key control points for caveola disassembly in response to an oxidizing environment.

## Introduction

Cellular membranes exhibit a complex composition, comprising thousands of distinct lipid species characterized by variations in their headgroups, chain lengths, and degrees of unsaturation. These lipid components are essential biomolecules for maintaining cellular architecture. However, the extensive diversity of membrane lipids poses challenges in understanding their precise roles and modes of action. A commonly accepted concept is the compartmentalization of the plasma membrane (PM) into specialized microdomains that control distinct biological functions. Therefore, unravelling the molecular roles of membrane lipids requires a detailed examination of their activities within these microdomains.

Caveolae are bulb-shaped lipid-protein microdomains that are highly abundant at the PM, particularly in specific cell types^1^. They possess a unique lipid composition and play crucial roles in a variety of biological processes such as endocytosis, signal transduction and lipid regulation^2,3^. As such, caveolae represent a distinct and valuable model system for studying the roles of specific lipid species in membrane function. The biogenesis of caveolae requires two families of proteins, caveolins and cavins^4–7^, and their interaction with membrane lipids. It is now clear that a comprehensive understanding of the role of the entire caveolar system, including its proteins, lipids, and architecture, is essential for elucidating the processes involved in the synthesis, maintenance, and function of caveolae.

Caveolins, small integral membrane proteins, work together with cytoplasmic proteins termed cavins, particularly Cavin1, to form caveolae. Other cavins (Cavin2, Cavin3 and muscle-specific Cavin4) associate with caveolae in a process dependent on Cavin1^8^. All cavins share conserved domains containing charged residues^5^ that allow them to interact with each other, Caveolin1 (CAV1) and lipids including phosphatidylserine (PS) and phosphatidylinositol 4,5 bisphosphate (PI(4,5)P2)^5,9–11^ through ‘fuzzy’ electrostatic interactions^4^. These findings suggest a model in which cavins, through their lipid-binding activities, establish a unique membrane lipid composition necessary for membrane-inserted CAV1 to sculpt the caveola membrane. Although the interactions of cavins, caveolins, and membrane lipids are individually of low affinity, they collectively, through a process termed ‘coincidence detection’, generate the characteristic caveolar membrane domain with distinct structure and lipid composition.

A crucial feature of this model, in which caveola formation relies on multiple low-affinity interactions, is that the domain is poised for a structural transition and can be rapidly disassembled. Since the lipid bilayer is relatively inelastic, caveola flattening can ensure the integrity of the PM in response to cell stretch or changes in cell shape. This process serves a protective role in various experiment models including cultured cells, tissue explants and animals^1,12,13^. Our recent studies show that caveolae are not only responsive to increased membrane tension but also to non-mechanical stimuli. Under UV or oxidative stress caveolae can partially disassemble and release cavins^14,15^. The released cavins can translocate to the cytosol and activate cell death pathways, specifically ferroptosis, in cells that fail to overcome oxidative insult as an evolutionarily conserved defence mechanism^14^. In this stress-sensing pathway, lipid peroxidation has been identified as an essential upstream mechanism for caveola disassembly^14^. It has been shown that Cavin1 dissociation from the cytoplasmic surface of caveolae and its interaction with downstream targets can be blocked or triggered by agents that respectively inhibit or stimulate lipid peroxidation^14^.

These recent discoveries suggest that our traditional view of caveolae needs re-evaluation and indicates that these membrane microdomains may act as general stress sensors capable of modulating the cellular signalling cascades through the co-function of caveolar structural proteins with their constituent lipids. The disassembly of caveolae therefore becomes a crucial aspect of this caveola signalling model. We hypothesise that unsaturated lipids prone to peroxidation are required for caveola formation and stability and that any changes in these lipids would alter caveolar structure and may induce caveola disassembly^2,16^. In this study we explored this novel aspect of membrane lipids and investigated the hypothesis that specific lipids serve as key regulators of both caveola formation and disassembly. We now demonstrate that specific unsaturated lipid species are required for caveola assembly and propose that this dependence provides a control point to initiate caveola disassembly in response to oxidative stress-induced lipid peroxidation.

## Results

### Targeted miniscreen for lipid enzymes involved in caveola formation

We first optimized confocal microscopy (CM), electron microscopy (EM) and biochemical methods for efficient assessment of caveola formation to allow medium throughput screening of key lipid enzymes. DHCR24 as a key enzyme involved in the synthesis of cholesterol (Chol)^17,18^, a known caveolar component^19^, was used as a positive control to optimize the methods for screening. We first confirmed that siRNA-mediated depletion of DHCR24 caused a significant decrease in caveolae in HeLa cells as determined by quantitative electron microscopy (EM) (Fig S1A). To facilitate higher throughput analyses required for screening lipid enzymes we optimized a high-resolution CM method to efficiently quantify caveolae. Immunolabeled endogenous Cavin1 (endo-Cavin1) was imaged in a focal plane (350 nm) at the basal cell membrane using CM with Airyscan detector (Fig 1A; see Method details). A punctate pattern indicates caveolar localization of Cavin1 (Fig 1B; S1B). Data was collected in an unbiased fashion by acquiring images from 5 randomly selected fields for each group (Fig 1A). Consistent with EM results, PM Cavin1 puncta (per cell) in DHCR24 knockdown (KD) cells were significantly reduced (Fig S1B-C), suggesting impaired caveola formation. These data demonstrated the effectiveness of our CM method in detecting and quantitating caveolae. In addition, we established a stable cell line expressing a newly developed nanobody against endo-Cavin1 with a C-terminal GFP tag (Fig S1D) as an independent parallel approach to track caveolae in live cells. Reduced GFP-nanobody signal at the PM indicated loss of caveolae in DHCR24 KD cells (Fig S1D). Consistent with partial disruption of caveolae upon KD of DHCR24, we also observed a reduction in levels of CAV1 and Cavin1 as judged by western blotting (Fig S1E).

**Figure 1.**
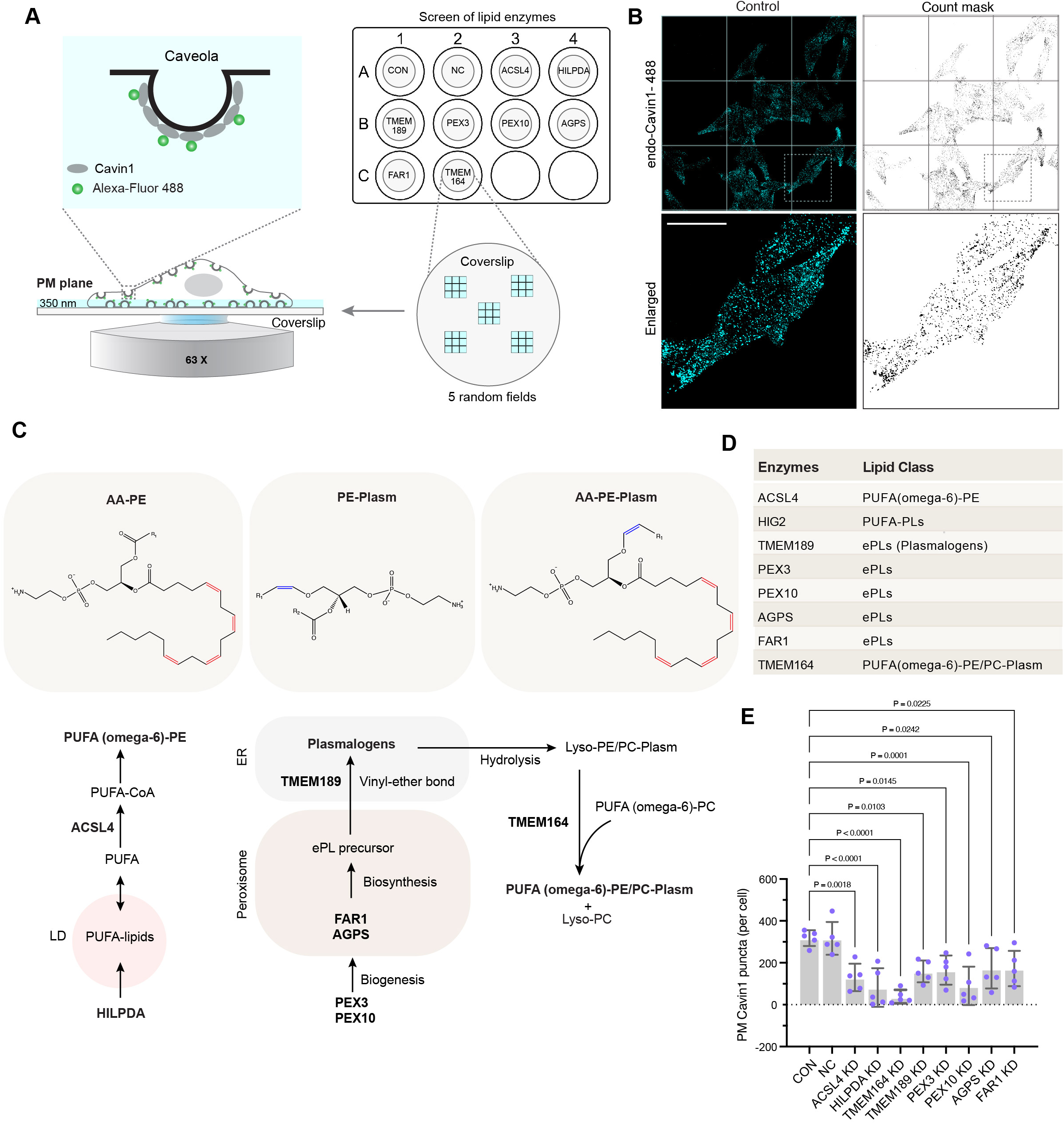
Screening of enzymes involved in caveola formation using quantitative CM. (A) A diagram illustrates quantitative CM-mediated assessment of the effect of knocking down target enzymes on caveola formation. HeLa cells seeded on coverslips in a 12-well plate were transfected with siRNAs targeting each enzyme in different wells. Scrambled siRNAs were transfected as the negative control (NC). Cells were fixed 48 h post transfection and endo-Cavin1 immunostained with Alexa-Fluor 488 at the basal membrane (350 nm plane) was imaged to indicate caveola abundance. (B) Confocal image showing PM Cavin1 puncta (cyan) in control group from 1 field (left panel). Right panel displays the mask of the counted puncta generated from image analysis. Scale bar = 10 μm. (C) A schematic overview of targeted enzymes and lipids. The top panel presents the structures of the targeted lipid classes, including PUFA-containing PLs (AA-PE), ePLs (PE-Plasm) and PUFA-containing ePLs (AA-PE-Plasm). Double bonds in the PUFA tail are highlighted in red, and the vinyl ether linkage in plasmalogens is highlighted in blue. The bottom diagram illustrates the localization and lipid-regulatory functions of the screened enzymes (highlighted in bold). Plasm: Plasmalogens. (D) A table displays the class of the lipid products synthesized by each screened enzyme. (E) Quantification of PM Cavin1 puncta per cell. Each dot represents the mean value of 1 random field (N = approx. 40 cells per field, 5 fields in total). Error bar = SD. Statistical analysis was performed using one-way ANOVA with Dunnett’s test.

Having established and characterized methods and systems for the evaluation of caveola abundance, we tested a small library of enzymes linked to lipid peroxidation and ferroptosis, including enzymes involved in the synthesis of polyunsaturated fatty acid (PUFA)-linked phospholipids (PLs)^20,21^ (ACSL4 and HILPDA) and ether lipids^22–24^ (PEX3, PEX10, AGPS, FAR1,TMEM189 and TMEM164) (see scheme in Fig 1C-D). Depletion of these enzymes can selectively block the synthesis of specific lipid species as characterized previously^20–24^. In this work we used siRNAs to deplete these enzymes in HeLa cells, creating an experimental system conductive to biochemical and cellular manipulations while minimizing the risk of compensatory changes commonly observed in knockout cells. Quantitative CM (see average Cavin1 intensity in images stacked from 5 fields in Fig S2A-B) revealed that depletion of each of the tested enzymes led to a significant reduction of caveolae (Fig 1E and S2C-D) with minimal effects on CAV1 and Cavin1 mRNA levels (Fig S2E-F). Together these screening results suggest that the tested pro-ferroptotic enzymes and their associated lipogenesis pathways contribute to caveola formation.

### ACSL4 plays a critical role in caveola formation in cells and in zebrafish

In view of the links between caveolae and ferroptosis^14^, we first focussed on ACSL4, a key pro-ferroptotic enzyme that esterifies long-chain PUFAs, particularly omega-6 (n-6) PUFAs arachidonic acid (AA, C20:4n-6) and adrenic acid (AdA, C22:4n-6)^25,26^, to promote their incorporation into phosphatidylethanolamine (PE)^21^. We further refined the quantitative CM method to measure caveolae per μm^2^ (see Method details) and complemented this with quantitative EM and *in vivo* studies. Treatment of cells with an ACSL4 inhibitor, rosiglitazone (ROSI)^21^ or KD of ACSL4 dramatically reduced caveola density (Fig 2A-G and S3A) with the remaining caveolae showing abnormal morphology (Fig 2F and S3B). Moreover, KD of HILPDA, proposed to regulate the supply of PUFAs for ACSL4 from LDs^27^, also reduced caveola abundance (Fig 2H and S4A-C).

**Figure 2.**
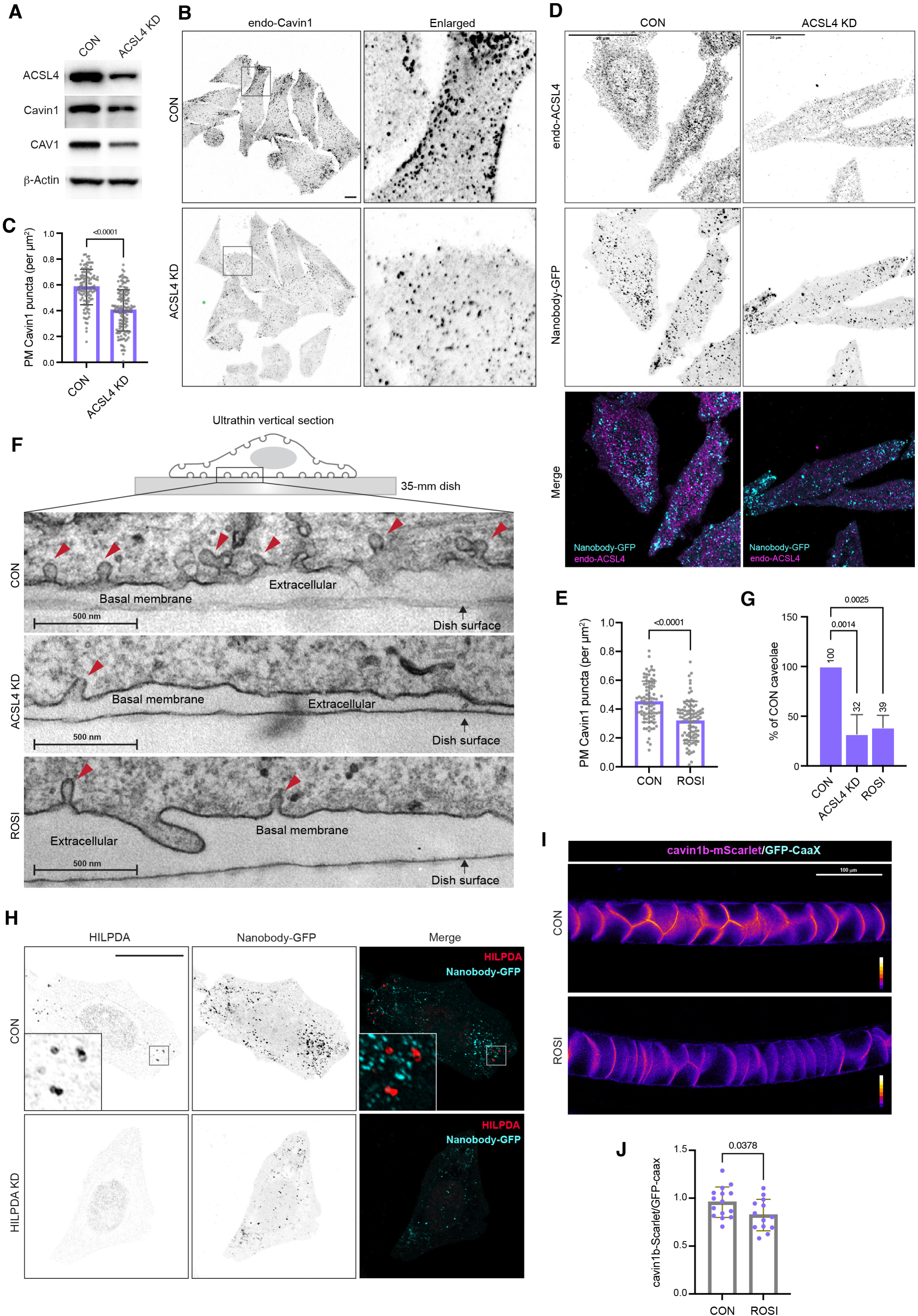
ACSL4 facilitates caveola formation in cells and in zebrafish. (A) Western blot showing the protein levels of ACSL4, Cavin1 and CAV1 in control (CON) and ACSL4 KD cells. (B) Inverted confocal images of PM Cavin1 in control and ACSL4 KD cells. Scale bar = 10 μm. (C) Quantification of PM Cavin1 puncta (per μm^2^/cell) in control and ACSL4 KD groups. Each dot represents an individual cell. N = approx. 120 cells for each group from 3 biological replicates. Error bar = SD. Statistical difference was assessed by two-tailed Student’s t-test. (D) Confocal images of nanobody against Cavin1 with a C-terminal tagged GFP (Nanobody-GFP, cyan) and immunostained ACSL4 (magenta) at the PM in control and ACSL4 KD cells. Scale bar = 20 μm. (E) Quantification showing decreased PM Cavin1 puncta (per μm^2^/cell) in ROSI-treated cells. Each dot represents an individual cell. N = approx. 90 cells for each group from 3 independent experiments. Error bar = SD. Statistical difference was assessed by two-tailed Student’s t-test. (F) EM images showing caveolae (red arrows) at the basal membranes of control, ACSL4 KD and ROSI-treated cells in an approx. 60 nm ultrathin vertical section. Scale bar = 500 nm. (G) EM analysis of caveola number relative to control group (%). Each dot represents the relative caveola count from different areas of the monolayers from two independent experiments. 25-30 images were quantified for each area. Error bar = SD. Statistical difference was assessed by one-way ANOVA with Dunnett’s test. (H) Confocal images of nanobody-GFP (cyan) and immunostained HILPDA (red) at the PM in control and HILPDA KD cells. Enlarged images showing the LD distribution of HILPDA in the control cells. Scale bar = 10 μm. (I) Calibrated ratiometric images (LUT “Fire”) showing the signal of cavin1b-mScarlet relative to PM marker GFP-CaaX in the notochord of control and ROSI-treated zebrafish (3 dpf). Scale bar = 100 μm. (J) Quantification of the intensity of cavin1b-mScarlet relative to GFP-CaaX in control and ROSI-treated zebrafish. Each dot represents an individual zebrafish (mixed clutch of 6 for each group). Statistical analysis was assessed by two-tailed Student’s t-test.

We next assessed the effect of ACSL4 inhibition on caveolae *in vivo* using a knock-in (KI) zebrafish line expressing cavin1b-mScarlet at endogenous levels (targeted insertion at the endogenous locus^28^). A strong fluorescent signal was observed in the notochord of 3 day post fertilization (dpf) embryos (Fig S3C), consistent with the high density of caveolae^12^. Quantitation of the membrane cavin1b-mScarlet signal relative to a stably expressed PM marker GFP-CaaX (Fig S3C) showed a significant reduction in membrane-associated cavin1b (see ROI selection in Method details and Fig S3D) in the notochord of the ROSI-treated zebrafish (Fig 2I-J), and caveola dysmorphology (Fig S3E-G). Together both *in cellulo* and *in vivo* data demonstrate that depletion or inhibition of ACSL4 disrupts caveolae and suggest a role of omega-6 PUFA-PE in caveola formation/stability.

### ACSL4 and TMEM189 facilitate caveola formation through independent pathways

We next turned to ether PLs (ePLs), a class of lipids that have been linked to both ferroptosis^22^ and caveolae^29^, characterized by the presence of either an ether linkage (plasmanyl-PLs) or a vinyl ether linkage (plasmenyl-PLs, also known as plasmalogens; Fig 1C) at the sn-1 position of the glycerol backbone. The synthesis of ePLs occurs through a peroxisome-dependent pathway^22^ or by the PUFA transferase, TMEM164, in a CoA-independent manner^24^. Our screening data (Fig 1E) and subsequent validation (Fig S4D-K) showed that KD of TMEM164 and the peroxisome-related enzymes including AGPS, FAR1, PEX3 and PEX10, led to a moderate but significant reduction of caveolae (Fig S4D-K). In contrast, KD of a non-lipid enzyme in peroxisomes, catalase (CAT), did not affect caveola abundance (Fig S4H-K). This suggests that ePLs are involved in caveola formation. Specifically, we demonstrated that TMEM189, the desaturase essential for introducing the vinyl ether bond during plasmalogen formation^30^, is critical for maintaining caveola abundance (Fig 1E; 3A-D), highlighting the specific role of plasmalogens, rather than general ePLs, in promoting caveola formation.

**Figure 3.**
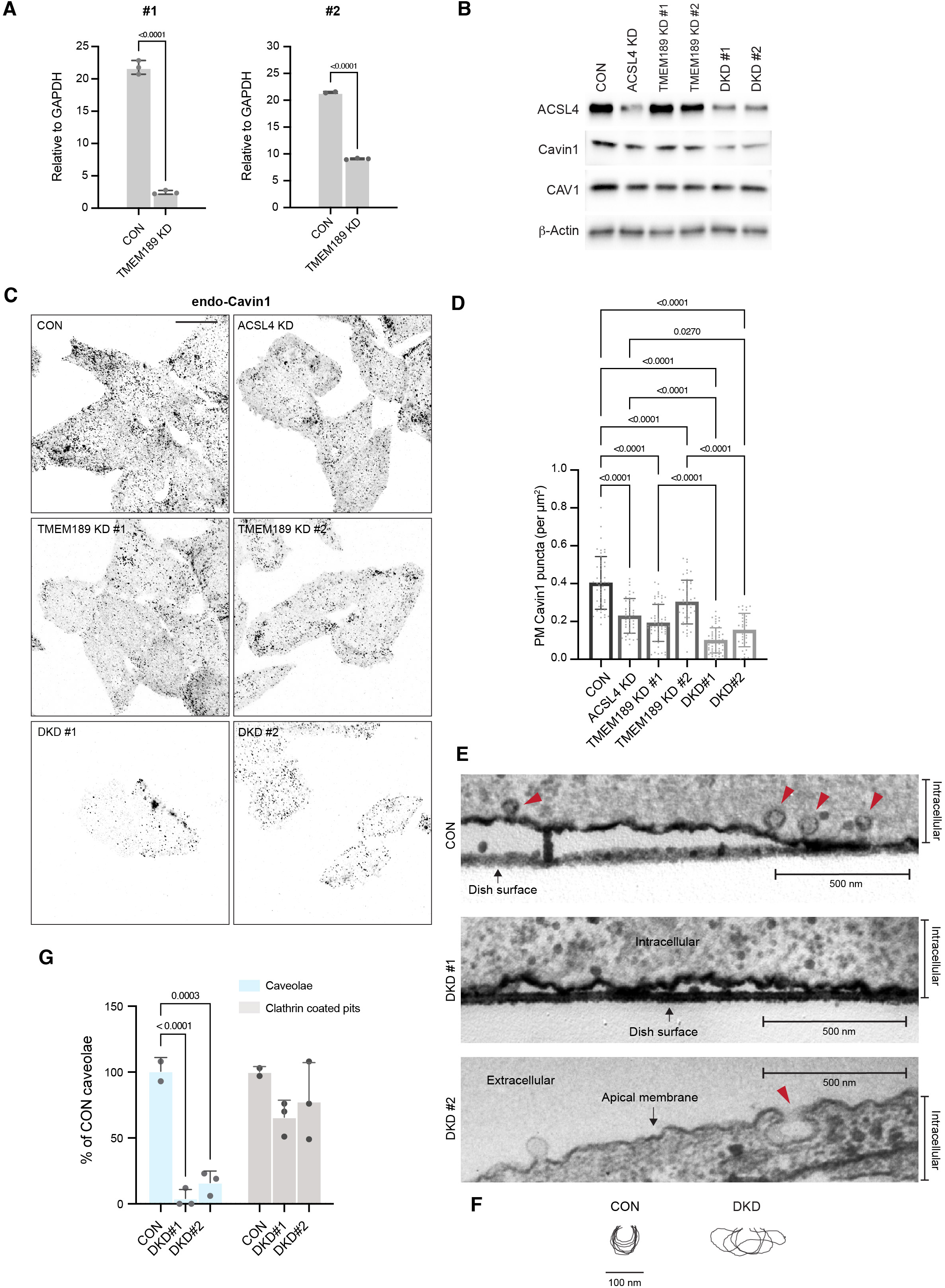
TMEM189 regulates caveola formation independently of ACSL4. (A) RT-PCR analysis of TMEM189 gene expression. The effect of two TMEM189 siRNAs were assessed. Each dot represents an independent biological replicate. Error bar = SD. Statistical difference was assessed by two-tailed Student’s t-test. (B) Western blot analysis of ACSL4, Cavin1 and CAV1 protein levels in the single KD and DKD cells. (C) Inverted confocal images of PM Cavin1 in the single KD and DKD cells. Scale bar = 10 μm. (D) Quantification of PM Cavin1 puncta (per μm^2^/cell) in (C). Each dot represents an individual cell. N = approx. 60 cells from 3 biological replicates. Error bar = SD. Statistical difference was assessed by one-way ANOVA with Tukey’s test. (E) EM images showing caveolae (red arrow) at the basal (control and DKD#1) or apical (DKD#2) membranes in an approx. 60 nm ultrathin vertical section. Scale bar = 500 nm. (F) Aggregated EM tracings of individual caveolae in control (N = 7) and DKD (N = 4) cells. (G) EM analysis of caveola number relative to control group (%). Clathrin coated pits were quantitated for each group as the negative control. Each dot represents the relative caveola count from different areas of the monolayer from two independent experiments. 25-30 images were quantified for each area. Error bar = SD. Statistical difference was assessed by two-way ANOVA with Dunnett’s test.

Given the significant impact of depleting ACSL4 or TMEM189 on reducing caveolae, and the secondary role of ACSL4 in synthesizing specific PUFA-containing PE plasmalogens^21^ that could also be formed by TMEM189^30^, we next investigated if these two enzymes promote caveola formation via a shared pathway involving plasmalogen synthesis. We observed that double KD (DKD) of ACSL4 and TMEM189 further reduced caveolae as compared to the single KD groups (Fig 3B-G); the number of caveolae was reduced 2 times more in the DKD cells (96.50 ± 14.35% reduction in DKD#1; 84.50 ± 14.35% reduction in DKD#2; Fig 3G) compared to the number in ACSL4 single KD cells (48% reduction; Fig 2G). Irregular morphology was also observed for many of the remaining caveolae in the DKD cells (Fig 3E-F). Together this suggests that ACSL4 and TMEM189 contribute to caveola formation through independent lipogenesis pathways, and that PUFA-PE (non-plasmalogen type) and plasmalogens both contribute to caveola formation.

Caveola biogenesis requires efficient CAV1 exit from the Golgi complex that is regulated by Chol and fatty acids (FAs)^31^ and CAV1 oligomerisation^32^. We evaluated if KD of ACSL4 and/or TMEM189 affects CAV1 exit from the Golgi complex to impair caveola biogenesis. CM revealed a decreased punctate PM CAV1 signal in both single KD and DKD cells (Fig S5A), consistent with the reduced PM Cavin1 in the KD cells (Fig 3C-D). However, there was no significant increase in CAV1 levels in the Golgi complex (Fig S5A). In contrast, KD of DHCR24 caused CAV1 retention in the Golgi complex (Fig S5B-C). This suggests that the exit of CAV1 from the Golgi complex was not significantly perturbed in cells lacking ACSL4 and/or TMEM189 and that ACSL4 and TMEM189 regulate caveola abundance through modulating the lipid composition of caveolae.

### Membrane PE species containing omega-6 PUFAs promote caveola formation

Our results implicate key regulatory enzymes in caveola formation/stability. We next investigated whether these effects are caused by loss of specific lipid species and can be rescued by their addition to cultured cells.

We focused on ACSL4 and observed that supplementation of its main omega-6 PUFA substrates, AA and AdA, could rescue caveolae in ACSL4-inhibited cells (Fig 4A-B). Activated AA/AdA (AA/AdA-CoA) by ACSL4 is preferentially incorporated into PE, rather than PC, to generate PE-AA/AdA as the major products^21^. We therefore also tested if addition of PE-AA (C18:0/20:4) could rescue caveolae in ACSL4-inhibited cells. The effect of PC-AA (C18:0/20:4) was also assessed as a negative control. Addition of PE-AA, but not PC-AA, could fully rescue the loss of caveolae in ACSL4-inhibited cells (Fig 4C-D) demonstrating that ACSL4 facilitates caveola formation through its key function in esterifying omega-6 PUFAs into PE.

**Figure 4.**
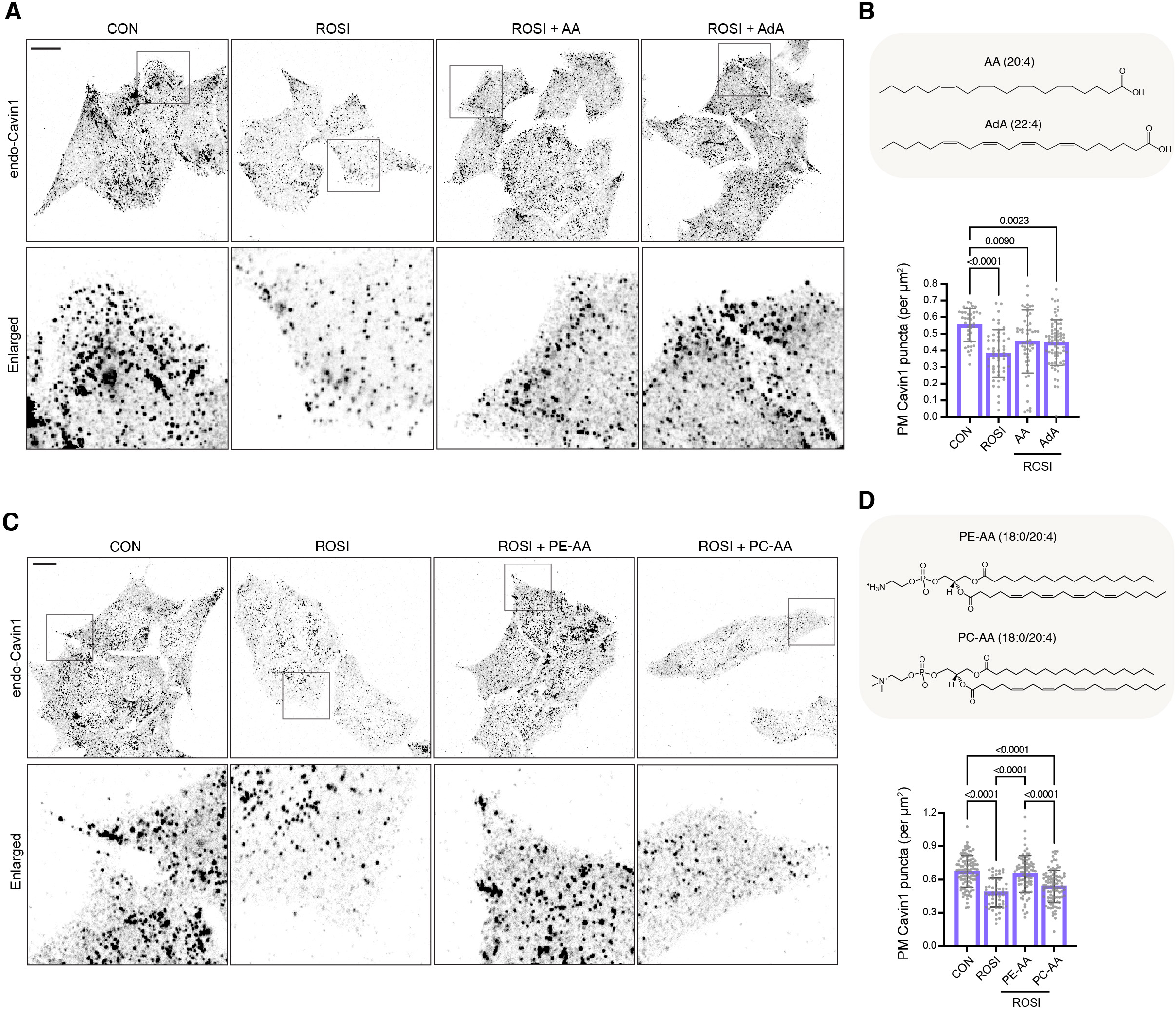
Supplementation of AA/AdA or PE-AA rescues caveolae in ACSL4-inhibited cells. (A) Inverted confocal images of PM Cavin1 in control or ROSI-treated cells with or without AA (15 μM) or AdA (15 μM) supplementation for 3 h. Scale bar = 10 μm. (B) The structures of supplemented omega-6 PUFAs and quantification of PM Cavin1 puncta in (A). Each dot represents an individual cell. N = approx. 60 cells for each group from 3 biological replicates. Error bar = SD. Statistical difference was assessed by one-way ANOVA with Tukey’s test. (C) Inverted confocal images of PM Cavin1 in control or ROSI-treated cells with or without SAPE (15 μM) or SAPC (15 μM) supplementation for 3 h. Scale bar = 10 μm. (D) The structures of supplemented PUFA-lipids and quantification of PM Cavin1 puncta in (C). Each dot represents an individual cell. N = approx. 90 cells for each group from 3 biological replicates. Error bar = SD. Statistical difference was assessed by one-way ANOVA with Tukey’s test.

The above studies rely on relatively long-term lipid supplementation (4 h in ACSL4-inhibited cells). To acutely modify PM lipids, we employed a recently described liposome system^33^. In this system, fusogenic liposomes with defined membrane lipid composition can immediately fuse with PM upon contact and thereby selectively and rapidly modify membrane lipid composition^33^. Fusogenic PE-AA-liposomes were added to untreated or ROSI-treated CAV1-mCherry-expressing HeLa cells^33^ for a short period (15 min) to insert PE-AA into the PM. Caveolae, as indicated by stably expressed CAV1-mCherry and immunostained Cavin1 at the PM, were significantly reduced in ACSL4-inhibited cells and could be rescued by acute treatment with fusogenic PE-AA-liposomes (Fig S6A-C). This suggests that membrane-incorporated PE containing omega-6 PUFAs plays a role in caveola formation.

### Displacement of membrane omega-6 PUFAs by omega-3 PUFAs and MUFA disrupts caveolae

Previous studies showed that dietary omega-3 (n-3) PUFAs could be incorporated into caveolae and displace omega-6 PUFAs in different classes of PLs, profoundly altering the lipid environment and the abundance of caveolae as indicated by CAV1 fractionation^34–37^. In addition, it has been shown that expression of MFSD2a - a lipid transporter for DHA uptake^38^ - could inhibit caveola formation^39^. We therefore tested if long-term supplementation of omega-3 PUFAs would impair caveola formation because of omega-6 PUFA displacement from caveolae.

We supplemented HeLa cells with different omega-3 PUFAs including docosahexaenoic acid (DHA, C22:6n-3) and eicosapentaenoic acid (EPA, C20:5n-3) (Fig 5A) in culture each day for 2 days and measured the effects on caveola formation. To minimize the effect of the FAs contained in the serum we decreased the serum concentration in the culture medium without affecting caveola formation (Fig 5B-D). Long-term incubation with DHA or EPA led to a reduction of caveolae (Fig 5B-D) whereas addition of AA did not affect caveola formation (Fig 5B-D). This demonstrated a negative regulatory effect of supplemented omega-3 PUFAs on caveola formation. However, acute modulation of PM lipids using fusogenic liposomes to directly insert PE-DHA (18:0/22:6) into the PM did not significantly affect caveola formation (Fig S6D). This suggests that the conversion of supplemented omega-3 PUFAs into esters and their subsequent incorporation into membrane lipids, replacing omega-6 PUFAs, causes disruption of caveola formation.

**Figure 5.**
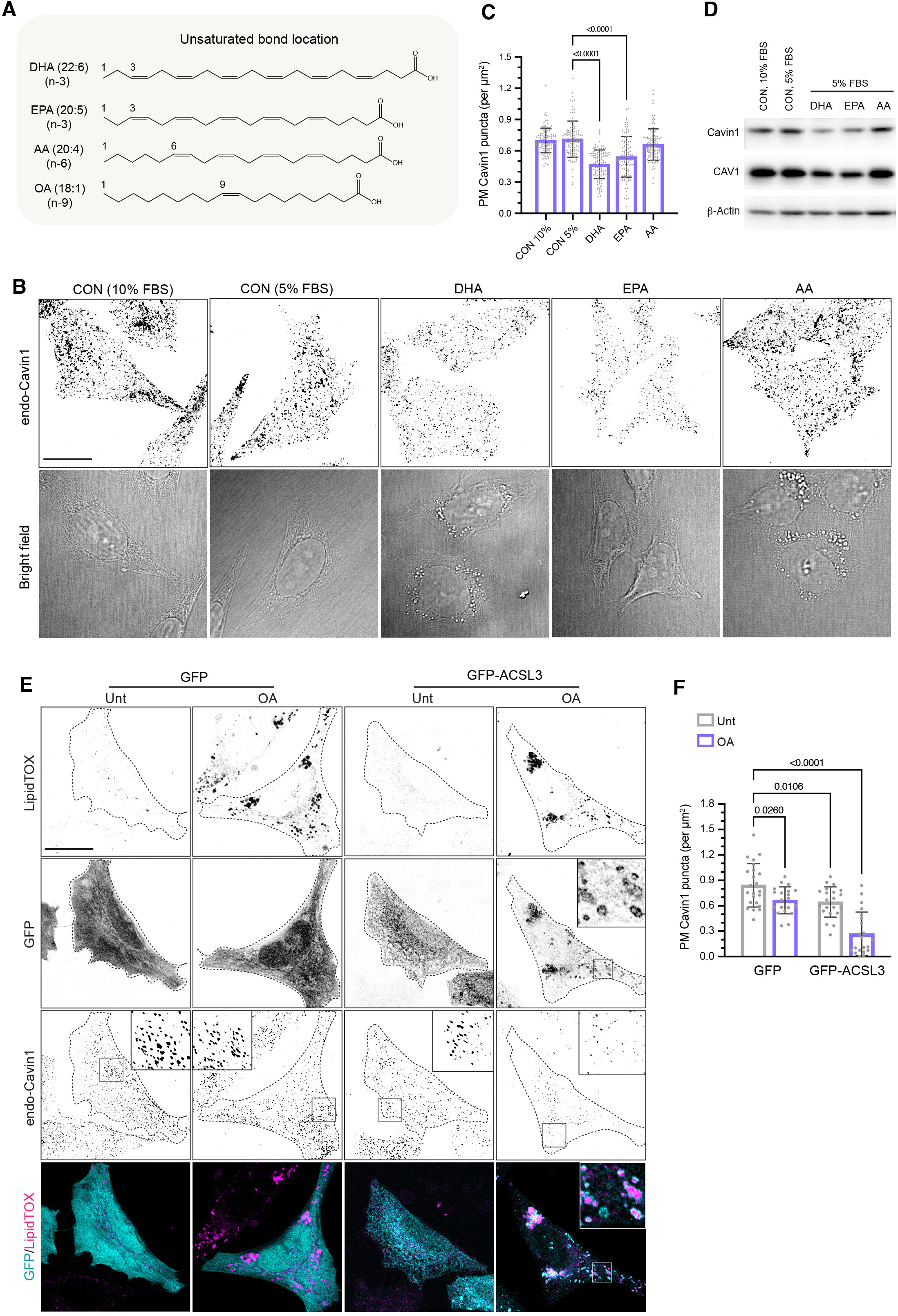
Long-term supplementation of omega-3 PUFAs and MUFA disrupts caveola formation. (A) The structures of supplemented omega-3 (DHA and EPA), omega-6 (AA) and MUFA (OA). (B) Inverted confocal images of PM Cavin1 (top panel) in cells with 48 h supplementation of different PUFAs (50 μM). Bright field images are displayed at the bottom. Scale bar = 10 μm. (C) Quantification of PM Cavin1 puncta (per μm^2^/cell) in (B). Each dot represents an individual cell. N = approx. 100 cells for each group from 3 biological replicates. Error bar = SD. Statistical difference was assessed by one-way ANOVA with Tukey’s test. (D) Western blot analysis of the protein expression of Cavin1 and CAV1 in PUFA supplemented cells. (E) Confocal images of PM Cavin1 (inverted) in cells expressing GFP-ACSL3 (cyan) with or without 24 h OA (50 μg/ml) treatment. GFP (cyan) was transfected as a negative control. Neutral lipids were stained with LipidTOX (magenta). GFP-ACSL3 shows LD localization in cells treated with OA (bottom right). Scale bar = 10 μm. (F) Quantification of PM Cavin1 puncta (per μm^2^/cell) in (E). Each dot represents an individual cell. N = approx. 20 cells for each group from 1 biological replicates. Error bar = SD. Statistical difference was assessed by one-way ANOVA with Tukey’s test.

In addition to dietary omega-3 PUFAs, exogenous monounsaturated fatty acids (MUFAs) can directly compete with omega-6 PUFAs for insertion into PM PLs^40^. It has been reported that activation of exogenous MUFAs by ACSL3, an ER- and LD-associated ACSL enzyme^41^, is required for their incorporation into membrane lipids^40^. We therefore evaluated the effect of supplementing MUFA oleic acid (OA, C18:1n-9) (Fig 5A) on caveola formation in cells with or without ACSL3 over-expression. CM revealed that over-expressed ACSL3 was mainly localized in the ER and could be recruited to LDs following OA treatment (Fig 5E), which is in agreement with previous findings^41^. Without ACSL3 overexpression, treatment with OA could reduce caveolae but with limited effect (17.8 ± 0.07% reduction; Fig 5F). ACSL3-overexpressing cells without OA treatment showed a similar small reduction in caveolae (19.7 ± 0.07% reduction; Fig 5F). The inhibitory effect of exogenous OA on caveola formation was significantly enhanced in cells with ACSL3 overexpression (57.35 ± 0.07% reduction; Fig 5F) suggesting that efficient MUFA activation and incorporation into the PM is required for its role in disrupting caveolae. These results demonstrate the essential role of omega-6 PUFAs in caveola formation and indicate that supplementation of FAs that displace membrane omega-6 PUFAs, such as omega-3 PUFAs and MUFAs, could negatively regulate caveola formation.

### Oxidized omega-6 PUFAs do not support caveola formation and trigger caveola disassembly

Taken together, the above results suggest that omega-6 PUFAs incorporated into membrane PLs such as PE through the action of ACSL4, work together with plasmalogens to maintain surface caveolae. As these lipids are major substrates for lipid peroxidation, and caveola disassembly can be induced upon lipid peroxidation^14^, we hypothesized that lipids containing oxidized omega-6 PUFA chains would be incompatible with caveola formation and so could form trigger points to drive caveola disassembly in response to oxidative stress.

The glutathione peroxidase 4 (GPX4) inhibitor, RSL3, which induces lipid peroxidation and ferroptosis^42^, can cause caveola disassembly^14^. It has been shown that doubly oxygenated hydroperoxyl-PE-AA/AdA (PE-AA/AdA-OOH) are the main oxidation products upon RSL3 treatment that executes ferroptosis^26^. We therefore compared the ability of oxidized PE-AA (PE-AA-OOH) and native PE-AA (Fig 6A) in promoting caveola formation and found that only native PE-AA could restore caveolae in ACSL4-inhibited cells (Fig 6B-C). These results showed a very specific requirement for PE-AA in caveola formation and suggest that PE that contains oxidized omega-6 PUFA chains loses the ability to promote caveola formation. To further test if oxidation of PE-AA/AdA could be the key regulatory mechanism for caveola disassembly, we compared the effect of RSL3-induced lipid peroxidation on caveola stability in control and ACSL4-inhibited cells. While RSL3 could effectively induce caveola disassembly in control cells, it did not affect caveola density in ACSL4-inhibited cells (Fig 6D-E). This shows that with reduced omega-6 PUFA containing PE at the PM, the remaining caveolae in ACSL4-inhibited cells become resistant to lipid peroxidation. Together these data suggest a critical role for omega-6 PUFA containing PE in caveola formation and as crucial control points to induce caveola disassembly upon their oxidation.

**Figure 6.**
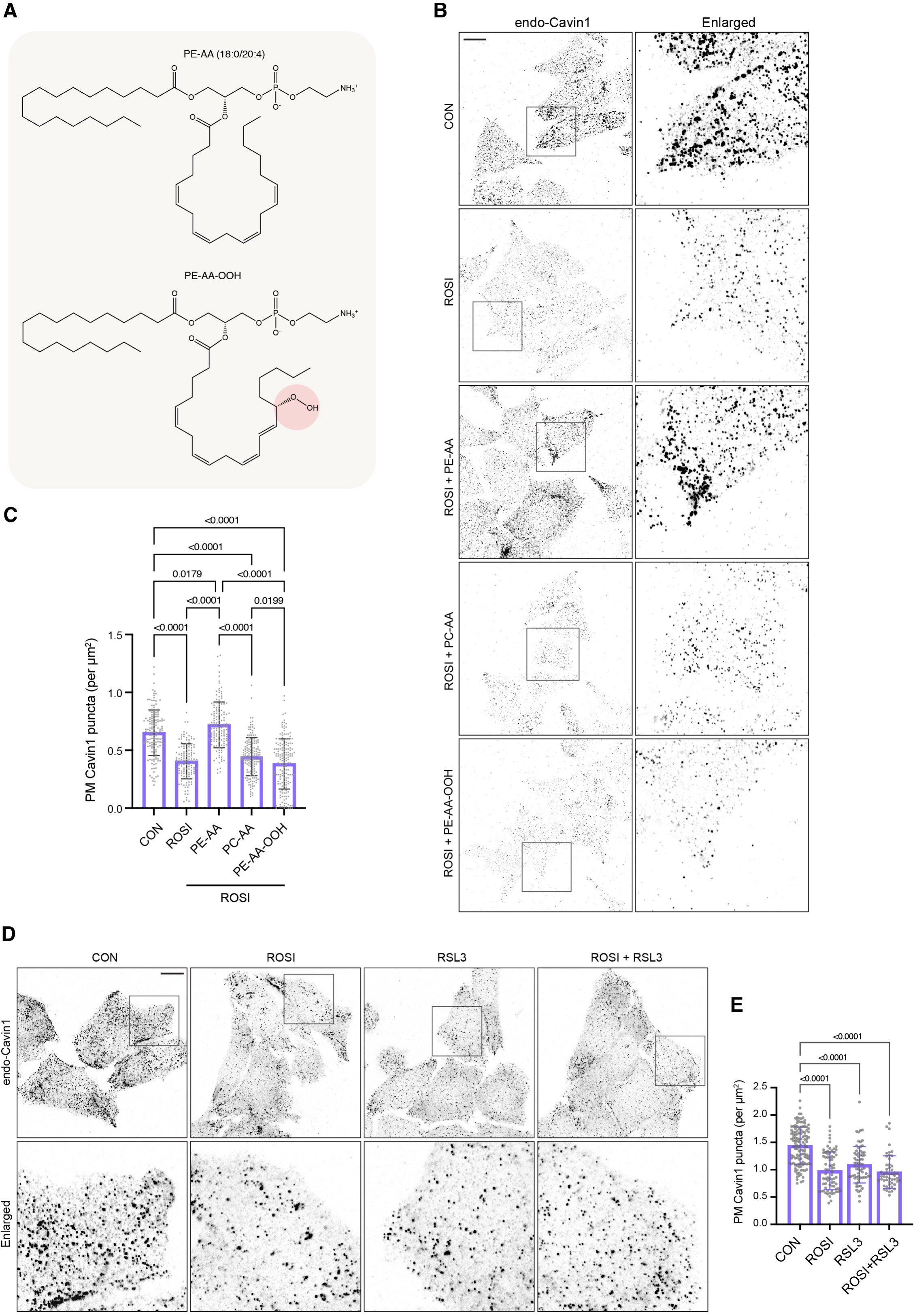
Oxidation of omega-6 attached to PE contributes to caveola disassembly. (A) The structures of native and oxidized PE-AA (PE-AA-OOH). (B) Inverted confocal images of PM Cavin1 in ROSI-treated cells with supplementation of native or oxidized lipids (15 μM). PC-AA was included as a negative control. Scale bar = 10 μm. (C) Quantification of PM Cavin1 puncta (per μm^2^/cell) in (B). Each dot represents an individual cell. N = approx. 150 cells for each group from 3 biological replicates. Error bar = SD. Statistical difference was assessed by one-way ANOVA with Tukey’s test. (D) Inverted confocal images of PM Cavin1 in control and ROSI-treated cells. RSL3 (10 μM) was added to the cells for 5 h post 24 h ROSI treatment. Scale bar = 10 μm. (E) Quantification of PM Cavin1 puncta (per μm^2^/cell) in (D). Each dot represents an individual cell. N = approx. 70 cells for each group from 3 biological replicates. Error bar = SD. Statistical difference was assessed by one-way ANOVA with Tukey’s test.

Given that electrostatic interactions between Cavin1 and specific PM lipids are fundamental for caveola formation^4,5,9–11^, we investigated if oxidation of omega-6 PUFAs disrupts Cavin1-membrane associations as a potential mechanism for caveola disassembly upon lipid peroxidation. To specifically evaluate Cavin1-membrane associations, we used HeLa *CAV1* knockout (KO) cells as a caveola-independent model. In this model, cytoplasmic Cavin1 can be recruited to the PM upon Chol addition bypassing the requirement for CAV1^4^. Without exogenous Chol, transiently expressed Cavin1-GFP diffusively localized in the cytoplasm in HeLa *CAV1* KO cells due to the absence of caveolae (Fig 7A). Upon Chol addition, Cavin1-GFP was recruited to the PM, forming punctate structures at low Chol concentration (4 μM) (Fig 7B and D) or membrane-associated tubules at high Chol concentration (20 μM) (Fig 7B and G), consistent with previous observations^4^. Cavin1-membrane associations formed rapidly 1 hour after Chol addition (Fig 7B) but were disrupted by RSL3 treatment (Fig 7C-D and 7G). Specifically, RSL3 treatment completely abolished Cavin1-membrane associations in 40% of cells with 4 μM Chol addition (Fig 7E) and significantly reduced the number of PM Cavin1 puncta (Fig 7F). At 20 μM Chol, both number and total area of PM-associated Cavin1 tubules per cell were significantly reduced after RSL3 treatment (Fig 7H-I). These data suggest that omega-6 PUFA oxidation during lipid peroxidation leads to Cavin1 dissociation from the PM, serving as a potential mechanism for caveola disassembly in an oxidizing environment.

**Figure 7.**
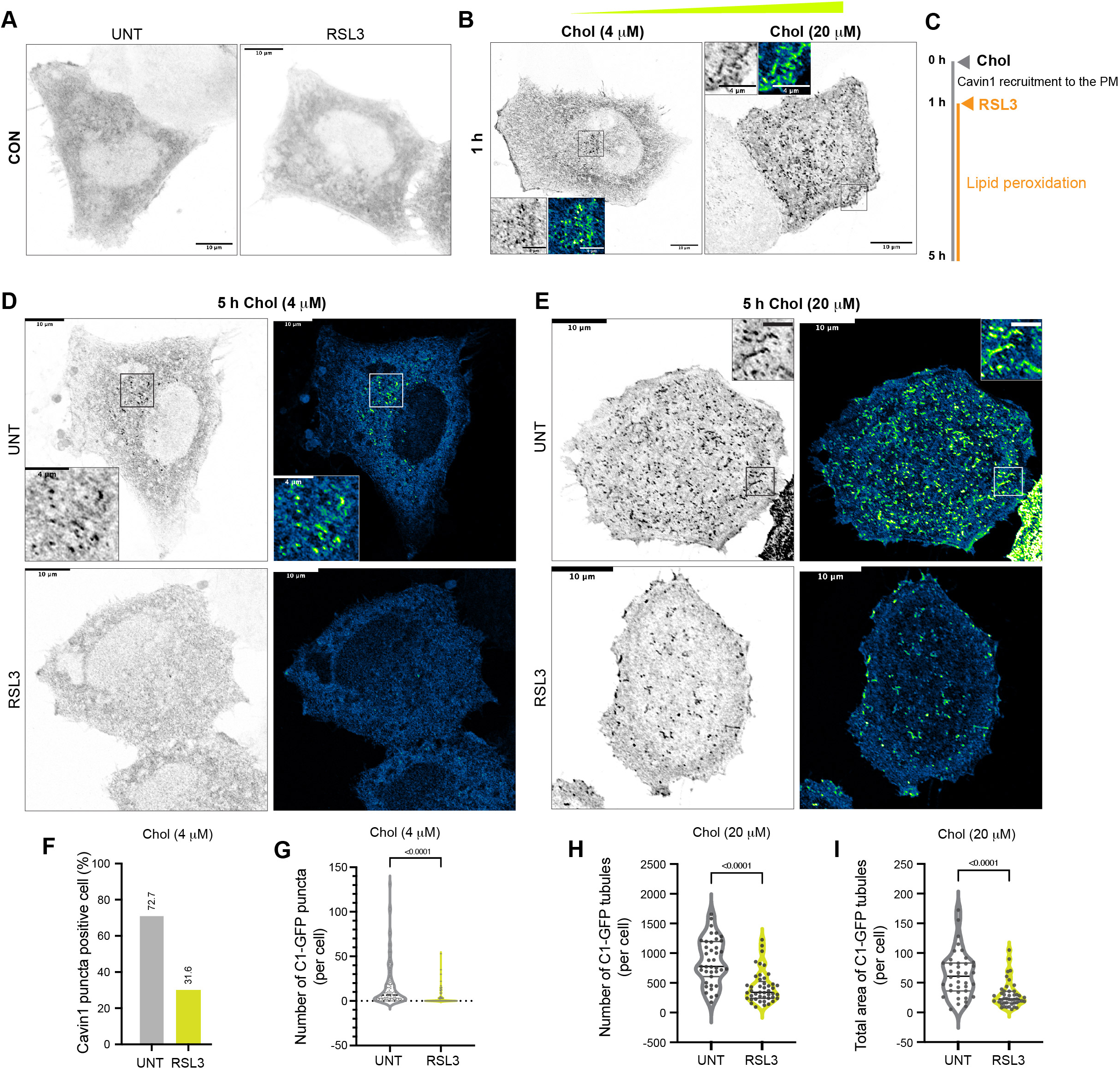
Lipid peroxidation induces Cavin1-membrane dissociation. (A) Inverted confocal images showing cytoplasmic localization of transiently expressed Cavin1-GFP in control HeLa *CAV1* KO cells. Scale bar = 10 μm. (B) Inverted single-plane (500 nm PM section) confocal images showing Cavin1-GFP recruitment to the PM 1 h after Chol addition at different concentrations. Lookup Table (LUT) “Green Fire Blue” was applied to the enlarged images. Scale bar = 10 μm; enlarged boxes scale bar = 4 μm. (C) Timeline of cell treatments with arrows indicating when Chol or RSL3 were added. (D-E) Inverted single-plane (500 nm PM section) confocal images (left panel) showing a reduction of membrane-associated Cavin1-GFP puncta at 4 μM Chol (D) or Cavin1-GFP tubules at 20 μM Chol (E) in RSL3-treated cells. LUT “Green Fire Blue” was applied to the same images (right panel). Scale bar = 10 μm; enlarged boxes scale bar = 4 μm. (F) Percentage of cells with membrane-associated Cavin1-GFP puncta at 4 μM Chol with and without RSL3 treatment. (G-H) The number of Cavin1-GFP puncta at 4 μM Chol (G) or Cavin1-GFP tubules at 20 μM Chol (H) per cell. Each dot represents an individual cell. Statistical difference was assessed by two-tailed Student’s t-test. (I) Total area (μm^2^) of C1-GFP tubules at 20 μM Chol per cell. Each dot represents an individual cell. Statistical difference was assessed by two-tailed Student’s t-test.

## Discussion

Caveolae have emerged as crucial cellular stress sensors. Here we uncover polyunsaturated membrane lipids, particularly omega-6 PUFAs incorporated into specific lipids, as key regulators of caveolar formation and sensitivity to oxidative stress. We propose a new paradigm in which caveolae have evolved to require specific lipid species, particularly those linked to ferroptotic signaling, for their formation and/or stability, providing a key control point for triggering their disassembly when these lipids become modified. The dependence of caveolae on these unsaturated lipids means that they form the ‘Achilles heel’ of the caveolae, a key vulnerability in an oxidizing environment allowing caveolae to be rapidly modulated, releasing cavins as signaling proteins into the cytosol where they can interact with a new set of intracellular proteins^14^. These results not only provide a new mechanistic understanding of how specific membrane lipids coordinate with membrane proteins to regulate cellular architecture and function, but also how changes in dietary lipids can have profound effects on cell and tissue function.

The identification of omega-6-linked PLs as specific lipids required for caveola formation that are also required to initiate caveola disassembly, raises the question of the site of action of these lipids. Our results suggest a role of omega-6 in caveola formation or stability at the PM, rather than during exocytic transport, as we observed no enrichment of CAV1 in the Golgi complex (as for example observed with Chol depletion upon DHCR24 knockdown; Fig S5) and a very rapid rescue of caveolae by direct insertion of PE-AA into the PM (within 15 minutes; Fig S6A-C). Downregulation of caveolar components observed with longer term knockdown of key lipid enzymes is consistent with disassembly of caveolae and degradation of their components as a secondary consequence, as observed in previous studies, although we cannot completely rule out direct effects of lipid loss on the stability of caveola proteins. Note that cytosolic Cavin1 is subject to proteasomal degradation^10^ and non-caveolar CAV1 to lysosomal degradation via VCP/p97^43^, and so would be predicted to be a consequence of caveolar disassembly. Nevertheless, our use of acute treatment with liposomes in rescue experiments argues for a very specific role in caveolar stability.

We propose that ePLs cooperate with PUFA-PLs to facilitate efficient caveola formation. Interestingly, through downregulation of TMEM189, our results implicated a specific role for plasmalogens. While all ePLs are susceptible to peroxidation due to their PUFA content, our results suggest that plasmalogen subtypes, characterized by their vinyl bonds at the sn-1 position, are particularly important for maintaining caveolae. The specific biophysical characteristics of plasmalogens may be important in this process but it is also intriguing that the vinyl bond of plasmalogens is particularly susceptible to oxidation^44,45^ and plasmalogens have been suggested to act as oxidative sinks^46^.

How might the properties of specific lipids influence caveola formation? Caveola formation involves a complex interplay between oligomeric caveolin discs^47^, Cavin1 oligomers^5^, and membrane lipids^11^. Previous work has shown a role for specific PS species in caveola formation, as well as reinforcing the complexity of the lipid environment generated independently but also synergistically by caveolins and cavins as they interact to form caveolae^11^. Lipid polyunsaturation is crucial for maintaining the structural and functional integrity of membranes. Unlike MUFAs and saturated fatty acids (SFAs) that are straight chained in membrane lipids, PUFAs can adopt various shapes from straight to highly bent, which increases membrane fluidity and elasticity^48^. PUFA-enriched membranes show decreased membrane bending stiffness, allowing for curvature formation^49^, which is essential for the generation of caveolar invaginations. Whether PUFAs play a general role in shaping caveolar membrane or a more specific role in facilitating the interdependent roles of caveolins and/or cavins must wait detailed molecular testing. However, our results showed that peroxidized PUFA-PE loses the ability to promote caveola formation and instead triggers caveola disassembly. By using a model cellular system lacking CAV1, we identified the disruption of Cavin1-membrane interactions upon lipid peroxidation as one mechanism that could potentially trigger disassembly. Molecular dynamics (MD) simulations have revealed that peroxidation alters lipid conformation and membrane biophysical properties in an oxidation site-dependent manner^50^. The effect of PUFA peroxidation in disrupting lipid-lipid packing becomes more pronounced as the peroxidation site moves towards the methyl end of the PUFA chain. For instance, peroxidation at C15 of the C20:4 PUFA tail, that corresponds to the lipid peroxide tested in this study (PE-AA-OOH; Fig 6A), has the most significant effect in reducing lipid tail packing order compared to C5/8/12-OOH, leading to membrane destabilization^50^. This indicates a potential role of the unique conformation created by the double bonds near the methyl end of PUFAs in stabilizing caveolae.

In addition, our data suggest that despite the biophysical similarities between PUFAs, omega-3 and omega-6 possess opposing roles in caveola formation. We revealed that omega-3 competes with omega-6 and so disrupts caveola formation. This is consistent with previous work linking dietary omega-3 to loss of CAV1 in caveolar fractions^37^. Due to the different locations of unsaturated bonds and degree of unsaturation in the acyl chain, omega-6 AA and omega-3 DHA show different conformational dynamics. AA has been proposed to form hairpin-shaped structures^51^ whereas tightly back-folded helical conformations are predicted to be most stable for DHA^52^. This difference could translate into distinct effects in shaping membranes with AA displaying an inverted conical shape supporting the formation of a positive curvature (bending towards the cytoplasmic side)^53,54^ and DHA a cone-shaped structure with a wider base than head that can induce a negative-curved membrane structure^54,55^.

Our results have revealed a striking parallel to studies of ferroptosis. Unsaturated FAs at the PM exert critical roles in ferroptosis execution. However, certain FAs can exhibit opposite effects on ferroptosis depending on the cell types or physiopathological conditions^56–58^. This suggests that the roles of unsaturated FAs in ferroptosis could not be solely attributed to their chemical features (e.g. degrees of unsaturation) but need to be evaluated within a comprehensive system that includes coordination with different cellular organelles and molecules. Our previous findings revealed that caveolae promote ferroptosis through inhibiting NRF2-mediated antioxidant response^14^. Therefore, comparing the roles of unsaturated FAs in ferroptosis and caveola formation may uncover potential downstream mechanisms that contribute to their distinct effects in ferroptosis. Omega-6 PUFAs promote ferroptosis^21,26^ and caveola formation, whereas MUFAs suppress both processes. This indicates that their differing effects on ferroptosis may be closely linked to their roles in caveola formation, which could subsequently affect caveola-mediated pro-ferroptotic signal. The role of omega-3 PUFAs in ferroptosis is more complex. Similar to omega-6 PUFAs, omega-3 PUFAs can induce ferroptosis as a reactant of peroxidation in acidic tumour environments^58^. However, they also protect neurons from ferroptosis by activating NRF2-mediated antioxidant response with unknown mechanisms in neurological disorders^56,57^. Considering the role of caveolae in inhibiting NRF2 activity^14^, we speculate that the anti-ferroptotic effect of omega-3 PUFAs may be related to their suppressive effects in caveola formation, as discovered in this study.

These results demonstrate the crucial role of specific lipid species in cellular processes. They also suggest mechanisms by which dietary lipids could influence cellular homeostasis, a subject of immense significance for population health. Omega-3 and omega-6 PUFAs as essential FAs that must be obtained from the diets, have garnered significant attention in dietary treatment and could influence lipid-regulated cellular processes such as inflammation and ferroptosis^58–60^. However, the limited understanding of the mechanisms involved hinders the application of dietary therapy. The studies presented here suggest that dietary changes in unsaturated lipids, which are known to modulate lipid composition throughout the entire body^61^, can differently affect the assembly and disassembly of caveolae. With caveolae being linked to vital homeostatic processes, such as the apical extrusion of tumour cells in epithelia^62^, the effects of dietary lipid changes could have long-term effects on health outcomes through modulation of caveolae and ferroptosis.

### Limitations of the Study

In this study we identify different roles of specific unsaturated lipids in caveola formation in a cell culture system and provide complementary evidence for this concept in a zebrafish model. We find that omega-6 PUFAs are crucial for caveola abundance whereas omega-3 PUFAs and MUFA antagonise their effects. However, we do not yet know how these unsaturated lipids modulate caveola abundance, whether it is through a direct effect on caveolar structure or more general effects on membrane structure. In addition, it is as yet unknown whether a similar lipid dependence would exist in all tissues or whether differences in lipid sensitivities can confer different properties in a cell-type specific manner.

## Acknowledgement

R.G.P. was supported by an Australian Research Council (ARC) Laureate Fellowship (FL210100107). This research is part of a project that has received funding from the European Research Council (ERC) under the European Union’s Seventh Framework Programme (FP7-2007-2013) (DRIMMS, Grant agreement No. 101071784). N.A. is supported by a Human Frontiers Science Program Grant (RGP011/2023). D.P.F. and J.L. were supported by an Australian NHMRC Investigator grant (2009551) and a grant from the Australian Research Council Centre of Excellence for Innovations in Peptide and Protein Science (CE200100012). The authors acknowledge the Microscopy Australia Research Facility at the Center for Microscopy and Microanalysis at The University of Queensland, the Biochemical Imaging Center (BICU) at Umeå University and the National Microscopy Infrastructure, NMI (VR-RFI 2016–00968). Confocal microscopy was performed at the Australian Cancer Research Foundation (ACRF)/Institute for Molecular Bioscience (IMB) Dynamic Imaging Facility for Cancer Biology with funding from the ACRF.

## Author Contributions

Conceptualization, Y.W. and R.G.P.; Methodology, Y.W., Y.-W.L., N.M., J.R., H.P.L., E.L., J.L., D.F. and T.E.H.; Formal Analysis, Y.W. and E.L.; Writing - Original Draft, Y.W. and R.G.P.; Writing - Review & Editing, Y.W., Y.-W.L., J.R., H.P.L., E.L., R.L., D.S.L., M.B., A.P., B.M.C., N.A., K.A.M., T.E.H. and R.G.P.; Supervision, Y.W. and R.G.P.; Funding Acquisition, N.A., D.P.F., A.P. and R.G.P.

## Declarations of Interests

The authors declare no competing interests.

## Inclusion and Diversity

We support inclusive, diverse, and equitable conduct of research.

## Supplemental Figure Titles and Legends

**Figure S1. DHCR24 is critical for caveola formation, Related to Figure 1**.

(A) EM analysis of caveola number relative to control group (%). Each dot represents an independent technical replicate. Error bar = SD. Statistical difference was assessed by two-tailed Student’s t-test.

(B) Confocal image showing PM Cavin1 puncta (cyan) from 1 random field (left panel) for each group. The mask of the counted puncta generated from image analysis were displayed. Scale bar = 10 μm.

(C) Quantification of PM Cavin1 puncta per cell. Each dot represents the mean value of 1 random field (N = approx. 40 cells per field). Error bar = SD. Statistical analysis was performed using two-tailed Student’s t-test.

(A) Confocal images of immunostained Cavin1 (magenta) and Nanobody-GFP (cyan) at the PM in control and DHCR24 KD cells. Scale bar = 10 μm.

(D) Western blot assays showing the protein levels of Cavin1 and CAV1 in control and DHCR24 KD cells.

**Figure S2. Quantitative CM-mediated evaluation of caveola abundance, Related to Figure 1**.

(A) A schematic illustrating the workflow of generating average Z stack projection of PM Cavin1 from 5 random fields for each group.

(B) Average Z stack projection showing an overall intensity of PM Cavin1 puncta from 5 random fields in each group. Scale bar = 50 μm.

(C-D) Representative images of PM Cavin1 (D) from a randomly selected region at the basal membrane (C) for each group. Scale bar = 4 μm.

(E-F) RT-PCR analysis of Cavin1 (E) and CAV1 (F) gene expression. Scrambled siRNAs were transfected as negative control (NC). Each dot represents a technical replicate. Error bar = SD. Statistical difference was assessed against NC by one-way ANOVA with Dunnett’s test.

**Figure S3. Caveola abundance in ACSL4-inhibited cells and zebrafish, Related to Figure 2 and STAR Method.**

(A) Confocal images of immunostained Cavin1 (magenta) and nanobody-GFP (cyan) at the PM in control and ROSI (10 μM, 24 h)-treated cells. Scale bar = 10 μm.

(B) Aggregated EM tracings of individual caveolae in control (N = 24), ACSL4 KD (N = 15) and ROSI-treated (N = 11) cells.

(C) Confocal images showing stably expressed cavin1b-mScarlet (magenta) and GFP-CaaX (cyan) in the notochord of the KI zebrafish line.

(D) Threshold-based generation of ROIs in zebrafish notochord. Gray scale cavin1b-mScarlet (top panel) was used for selecting membrane regions of zebrafish notochord as ROIs. The Moments threshold was applied to cavin1-mScarlet channel to generate binary image which was then delineated as ROIs. Bottom panel displays the mask of selected ROIs.

(E) EM images showing caveolar morphology in the notochord of control and ROSI-treated zebrafish in an ultrathin (approx. 60 nm) section. Scale bar = 1 μm.

(F) Aggregated EM tracings of individual caveolae in control (N = 98) and ROSI-treated (N = 131) zebrafish.

(G) Quantification of the lengths in pixels of the 2D tracings of caveolae presented in (F). Error bar = SD. Statistical analysis was performed using two-tailed Student’s t-test.

**Figure S4. The effects of HILPDA and ether lipid enzymes on caveolae, Related to Figure 2 and 3**.

(A-C) The effect of KD of HILPDA on the protein expression levels of Cavin1 and CAV1 (A) and on PM Cavin1 density (B-C). Each dot in (C) represents an individual cell (N = approx. 90 cells for each group from 3 biological replicates; Error bar = SD). Statistical analysis was performed using two-tailed Student’s t-test. Scale bar = 10 μm.

(D-F) The effect of KD of TMEM164 on the protein expression levels of Cavin1 and CAV1 (E) and on PM Cavin1 density (F-G). Enlarged region in (B) displays LD-localized HILPDA in the control cells. Each dot in (G) represents an individual cell (N = approx. 60 cells for each group from 3 biological replicates; Error bar = SD). Statistical analysis was performed using two-tailed Student’s t-test (G). Scale bar = 10 μm.

(G) Confocal images of nanobody-GFP (cyan) and immunostained TMEM164 (red) at the PM in control and TMEM164 KD cells. Scale bar = 10 μm.

(H-I) Western blot assays showing the effects of knocking down different peroxisomal enzymes on the protein expression levels of Cavin1 and CAV1 (H). The knockdown efficiency for AGPS, FAR1 and CAT was assessed (I).

(J-K) Confocal images (J) and quantification (K) showing the effects of knocking down different peroxisomal enzymes on PM Cavin1 density. Each dot in (L) represents an individual cell (N = approx. 100 cells for each group from 3 biological replicates; Error bar = SD). Statistical analysis was performed using one-way ANOVA with Dunnett’s test. Scale bar = 10 μm.

**Figure S5. Distribution of CAV1 in cells depleted with different enzymes, Related to Figure 3**.

(A) Inverted confocal images of CAV1 distribution in control, ACSL4 KD, TMEM189 KD and ACSL4/TMEM189 double KD (DKD) cells. Whole-cell (top panel), PM (middle panel) and intracellular CAV1 signals (bottom panel) are displayed. The whole-cell images were generated from z-stack images using maximum intensity projection. Scale bar = 10 μm.

(B-C) Confocal images showing endogenous CAV1 (cyan) and GM130 (magenta) in control (B) and DHCR24 KD (C) cells. Whole-cell (top 2 panels), PM (middle panel) and intracellular signals (bottom 2 panels) are displayed for both proteins. The whole-cell images were generated from z-stack images using maximum intensity projection. Enlarged images show the localization of CAV1 in the Golgi complex as indicated by GM130. Scale bar = 10 μm.

**Figure S6. Acute modification on membrane lipid content by fusogenic liposomes, Related to Figure 4 and 5**.

(A) Total internal reflection fluorescence (TIRF) images of stably expressed CAV1-mCherry (magenta) and immunostained endo-Cavin1 (cyan) at the PM in control and ROSI-treated cells. Fusogenic PE-AA-liposomes were added to the cells for 15 min prior to cell fixation. Scale bar = 10 μm.

(B-C) Quantification of PM CAV1-mCherry (CAV1-mCh) (B) and endo-Cavin1 (C) puncta in (A). Each dot in represents an individual cell (N = approx. 30 cells for each group from 1 biological replicate; Error bar = SD). Statistical analysis was performed using one-way ANOVA with Dunnett’s test.

(D) Relative change (%) of PM CAV1-mCherry (CAV1-mCh) puncta (per μm^2^/cell) after 15 min treatment of fusogenic PE-AA- or PE-DHA-liposomes. PM CAV1-mCherry puncta in live cells were imaged using TIRF at 0 min and 15 min after addition of fusogenic liposomes. Each dot in represents an individual cell (N = approx. 20 cells for each group from 1 biological replicate; Error bar = SD). Statistical analysis was performed using one-way ANOVA with Dunnett’s test.

## STAR Methods

### Resource availability

#### Lead contact

- Further information and requests for resources and reagents should be directed to and will be fulfilled by the Lead Contact, Robert G. Parton.

#### Materials availability

- Stable Cavin1-nanobody-GFP HeLa cell line, HeLa *CAV1* KO cell line and cavin1b-mScarlet KI zebrafish generated in this study are available upon request from the Lead Contact.
- The identifier numbers for plasmids used in this study have been listed in the Key Resource Table. The details of generation and validation of nanobody against Cavin1 will be reported elsewhere, and its sequence is available on request.
- This study did not generate new unique reagents.

#### Data and code availability

- Raw data of western blot analysis from Figures 2, 3, 5, S1 and S4 have been deposited on Mendeley Data repository and are publicly available. The DOI is listed in the Key Resource Table.
- This study did not use any unreported custom code or mathematical algorithm that is deemed central to the conclusion.
- Any additional information required to reanalyze the data reported in this work paper is available from the Lead Contact upon request.

### Experimental model and subject details

#### Zebrafish (Danio rerio)

Zebrafish were raised and maintained according to institutional guidelines (Techniplast recirculating system, 14-h light/ 10-h dark cycle, The University of Queensland). Adults (90 dpf above) were housed in 3 or 8 L tanks with flow at 28.5 °C, late-larval to juvenile stage zebrafish (6 dpf to 45 dpf) were housed in 1 L tanks with flow at 28.5 °C and embryos (up to 5 dpf) were housed in 8 cm Petri dishes in standard E3 media (5 mM NaCl, 0.17 mM KCl, 0.33 mM CaCl_2_ and 0.33 mM MgSO_4_) (Westerfield, 2000) at 28.5 °C (incubated in the dark). All experiments were performed in accordance with relevant guidelines and regulation with the approval of the University of Queensland (UQ) Animal Ethics Committee (Molecular Biosciences Animal Ethics Committee) and UQ Biosafety Committee. The following zebrafish strains were used in this study: wild-type (TAB; Tuebingen/AB) derived from the Zebrafish International Resource Centre Oregon, [*TgKI(cavin1b-mScarlet)^pd126^*] (this study), *Bact2-EGFP-CaaX^pc10Tg^*^63^. Zebrafish of developmental stages up to 15 dpf are prior to specific sex determination^64^ and the developmental stages are specifically stated in corresponding figure legends. All zebrafish used in this study were healthy, not involved in previous procedures and drug or test naïve. All zebrafish of the same clutch, and sex or developmental stage were randomly allocated into experimental groups.

#### Cell lines

The HeLa cell line was purchased from the American Tissue Culture Collection (ATCC) and grown in Dulbecco’s modified Eagle’s medium (DMEM) with 10 % (vol/vol) fetal bovine serum (FBS) at 37 °C in a humidified atmosphere containing 5% CO_2_. The HeLa Flp-In T-REx CAV1-mCherry cells were maintained in DMEM with 10 % (vol/vol) FBS and 5 μg/ml blasticidin S HCl (Thermo Fisher Scientific) for plasmid selection as described previously^33^. In addition, regular mycoplasma test was performed in our laboratory for cell line authentication and all cell lines used tested negative as assessed by MycoAlert Mycoplasma Detection Kit (Lonza Australia).

### Method details

#### Animal handling and reagents

Zebrafish embryos up to 7 dpf were raised and handled in standard E3 media during experiments (5 mM NaCl, 0.17 mM KCl, 0.33 mM CaCl2, 0.33 mM MgSO4). All post-embryonic zebrafish measurements were carried out between tanks of similar population densities and conditions. All reagents were obtained from Sigma-Aldrich unless otherwise specified.

#### Generation of cavin1b-mScarlet KI zebrafish

The cavin1b-mScarlet KI [*TgKI(cavin1b-mScarlet)^pd126^*] zebrafish line was generated using previously described methodology^28^. A gene fragment spanning from the end of intron 1 to the last coding sequence codon of exon 2 was cloned into pUC19-TgKI-MCS-mScarlet-MCS-polyA^28^ using In-Fusion seamless cloning (Takara Bio). Two gRNAs were synthesized as described using the HiScribe T7 Quick High Yield RNA Synthesis Kit (NEB) and purified with the Monarch RNA cleanup kit (NEB). The composition of the final injection mix: PCR donor (7.5 nM), gRNA (50 ng/µl), Cas9-NLS protein (5 µg/µl) (PNA Bio CP-02), Phenol Red (0.05%).

#### Lentivirus production and transduction

Lentiviral particles were produced as previously described^65^ with minor modifications. Briefly, HEK-293T cells were grown in DMEM supplemented with 10% FBS, 1% non-essential amino acids and 2mM L-glutamine at 37°C with 5% CO_2_. To produce lentiviral particles, plasmid for GFP-tagged nanobody against Cavin1 (sequence available upon request, see details below) and packaging vectors (pMDLg/pRRE, pRSV-Rev and pMD2.G^66^; see Addgene IDs in the Key Resource Table) were transfected into HEK-293T cells using Lipofectamine 2000 and growth media was replaced 24 h post-transfection. After another 48 h, virus-containing growth medium was collected, centrifuged at 2000 rpm for 7 mins and filtered through a 0.45 μm syringe-driven filter to remove cell debris. Virus-containing media was concentrated using an Amicon® Ultra-15 centrifugal filter unit. Concentrated virus-containing media was used for transduction of HeLa cells in standard growth media containing 10 μg/ml Polybrene Infection/Transfection reagent. GFP-positive HeLa cells were selected by flow cytometry (BD FACSAria Cell Sorter, Flow Cytometry Facility, Queensland Brain Institute, UQ).

#### Lentiviral transfer plasmid

To facilitate straightforward production of transfer plasmids we constructed a lentiviral gateway destination vector compatible with existing multisite gateway entry clone libraries. The multiple cloning site of pLL5^67^ was replaced with a multisite gateway cassette containing the chloramphenicol and ccdb resistance open frames flanked by AttR1 and AttR3 sites (Genscript). The resultant plasmid was termed pDEST-pLL5Gv2. The transfer plasmid pLL5Gv2-Cavin1_B7[VHH]-EGFP was made by recombining this destination vector with the entry clones pME-VHH[Cavin1_B7], p3E-EGFP^68^ (see Addgene IDs in the Key Resource Table).

#### Generation of nanobody against Cavin1

The alpaca-derived nanobody against Cavin1 was generated in collaboration with Wai-Hong Tham (Walter and Eliza Hall Institute, Melbourne Australia). The details of its generation and validation will be reported elsewhere, and its sequence is available on request. The sequence was subsequently cloned into pEGFP-N1 for expression with a C-terminal GFP-tag for experiments reported here.

#### Generation of HeLa CAV1 KO cells

Human *CAV1* gene was targeted by a CRISPR-Cas9 system^69^. *CAV1*-targeting sgRNA sequence was inserted into the BbsI site of PX459 pSpCas9(BB)-2A-Puro V2.0 vector^69^ using a synthesised oligonucleotide. Correctly inserted *CAV1* sgRNA sequence was confirmed by Sanger sequencing with hU6 forward primer. HeLa cells were then transfected using Lipofectamine 3000. 48 h post transfection, HeLa *CAV1* KO cells were assessed using western blot to confirm KO of CAV1 protein.

#### Immunofluorescence

Cells were plated onto glass coverslips at 70% confluence and then fixed in 4% (vol/vol) paraformaldehyde in PBS for 15 min at room temperature (RT) after the indicated treatments. Coverslips were washed three times in PBS and permeabilized in 0.1% (vol/vol) Triton X-100 in PBS for 5 min and blocked in 1% (vol/vol) BSA in PBS for 30 min at RT. Primary antibodies were diluted in 1% BSA/PBS solution at optimized concentrations and incubated with coverslips for 1 h at RT. Diluted secondary antibodies (1:500 in 1% BSA/PBS) conjugated with fluorescent dyes were later added onto coverslips and incubated for 1 h. For neutral lipid staining, diluted LipidTOX (Thermo Fisher Scientific) (1:200) was added on coverslips for 30 min after incubation with secondary antibodies. Coverslips were washed in PBS for three times and mounted in Mowiol in 0.2 M Tris-HCl pH 8.5.

#### Cell treatment

For FA or lipid supplementation, cells were seeded on a coverslip or plate at 70% confluence and cultured in DMEM with reduced FBS (5%) for 18 h. On the following day, FAs (AA, AdA, DHA or EPA) or lipids (PE-AA, PE-AA-OOH or PC-AA) was diluted to indicated concentrations in DMEM containing 5% FBS and 2% fatty acid free bovine serum albumin (BSA) and then added to the cells for the specified duration as per the legends. For long-term FA supplementation, PUFAs (DHA, EPA and AA) were replaced every 24 h. For ACSL4 inhibition, cells were treated with 10 μM Rosiglitazone (ROSI) (Sigma-Aldrich) for 24 h prior to FA or lipid supplementation.

Chol addition experiments were performed in HeLa *CAV1* KO cells transiently expressing GFP tagged Cavin1^70^ using previously described methodology^4^. Water-soluble analogue of Chol (Sigma-Aldrich) was diluted to indicated concentrations in normal DMEM culture medium containing 10% FBS and then added to the cells for the specified duration as per the legends.

#### Confocal microscopy and image analysis on cells

The basal membranes of fixed cells were imaged by Zeiss LSM 900 confocal microscope with Airyscan 2 detector and a 63 X oil lens. Airyscan detector was set to SR mode via Zeiss Zen interface for high-resolution imaging to achieve a 350 nm axial resolution. Super resolution processing was applied to acquired images using Zen. For the miniscreen experiment, five random fields on each coverslip were selected (approx. positions as shown in Fig 1A) using the multi-position function in Zen with a 3 X 3 tiled imaging method.

The basal membranes of fusogenic liposome-treated live or fixed cells were imaged using TIRF on Zeiss Spinning Disk confocal microscope controlled by the ZEN interface with an Axio Observer.Z1 inverted microscope, equipped with an EMCCD camera iXonUltra from ANDOR (100 X lens) as described previously^33^.

Image analysis was performed using ImageJ 2.1^71^. For cell size measurement (μm^2^), the Li threshold was applied to grayscale images to distinguish the entire area for each cell. Cell area was then measured using ‘Analyze particles’ with size range manually optimized for each experiment based on the image resolution. Each cell area was saved as region of interest (ROI). The same raw images were then subjected to puncta quantification. Background subtraction (> 50 pixels) was performed, followed by the MaxEntropy threshold to the images. Puncta within each cell was counted using ‘Analyze particles’ within each ROI saved from the cell size measurement step. To account for cell size variations, the puncta count was normalized by dividing by the cell size. The same parameter settings were consistently and automatedly applied across all groups within each experiment.

#### Live confocal imaging and image analysis on zebrafish embryos

Live confocal microscopy on zebrafish was performed as previously described^14^. For ACSL4 inhibition, 10 hpf embryos were first dechorionated by incubation in pronase (0.05 mg/mL) for 2 h at RT. At 12 hpf, embryos were treated with 20 μM ROSI [in a solution of 1 % DMSO and 0.2 mM phenylthiourea (PTU) in standard E3 media (5 mM NaCl, 0.17 mM KCl, 0.33 mM CaCl_2_, 0.33 mM MgSO_4_)] and incubated up until 3 dpf. The same solution without ROSI was used in the control group. At 2 dpf, cavin1b-mScarlet positive zebrafish embryos were screened. Prior to imaging, zebrafish (3 dpf) were anesthetized in tricaine solution in E3, mounted in 1% low melting point (LMP) agarose on MatTek glass bottom dishes in a lateral view unless otherwise stated (anterior to the left, posterior to the right) and submerged in tricaine solution. The embryos were submerged in tricaine solution in E3 throughout all imaging periods. For live confocal images of WT and cavin1a/1b DKO zebrafish, embryos were incubated at 28°C and mounted in LMP agarose as above and imaged under a Zeiss LSM880 confocal microscope.

To minimize bias a blind image analysis was performed. The image file names were encrypted by the image collector (Y.-W.L.) and then blindly analyzed by a different individual (Y.W.) who was unaware of the treatment groups. ImageJ 2.1^71^ was used for image analysis. The Moments threshold was applied to grayscale cavin1b-mScarlet channel to generate binary image. This was followed by “Analyze Particles” (size = 10-infinity pixel) for identifying the membrane regions of the notochord, which were saved and combined into a single ROI for each image (see ROI generation in untreated zebrafish in Fig S3D). The cavin1b-mScarlet intensity within the ROI was measured and normalized to the GFP-CaaX intensity within the same region. Encrypted names were revealed only after image analysis to allocate quantitated data to their respective groups. The same parameter settings were consistently and automatedly applied across all images.

#### Electron microscopy

EM on cells^15^ and zebrafish^12,14^ was performed as previously described. Briefly, cells and zebrafish tissue were initially fixed in 2.5% glutaraldehyde, then post fixed and contrasted in a series of immersions: 1.5% potassium ferricyanide and 2% osmium tetroxide, 1% thiocarbohydrazide, 2% osmium tetroxide, 2% uranyl acetate and finally 0.06% lead nitrate using a Pelco Biowave at 80W under vacuum for 3min each. Samples then underwent serial dehydration in increasing concentrations of ethanol, before serial infiltration with increasing concentrations of LX112 resin for cells and Procure 812 resin for zebrafish tissue. Ultrathin sections (approx. 60 nm) were obtained using a Leica UC64 ultramicrotome and imaged on a Jeol JEM-1011 at 80kV.

#### RNA preparation, reverse-transcription and quantitative real-time PCR

Total RNA was extracted using QIAGEN RNA isolation kit following the manufacturer’s instructions and measured using a NanoDrop spectrophotometer (ThermoFisher Scientific). Two-step reverse transcription was conducted to access single strand cDNA using SuperscriptIII Reverse Transcriptase (ThermoFisher Scientific) as per manufacturer’s instruction. SYBR-Green or Taqman PCR Mater Mix was utilized for real-time PCR using a ViiA7 Real-time PCR system (ThermoFisher Scientific). The sequences of oligonucleotide primers for real-time PCR are listed in Key Resources Table. Gene expression was analyzed using the delta-delta Ct method as previously described^72^.

#### Cell transfection

Cells were plated at 70% confluence for DNA transfection, or at 50% confluence for siRNA transfection on a plate or on coverslips. On the following day, DNA constructs or siRNA were transfected using Lipofectamine 3000 (DNA) or 2000 (siRNA) kit (ThermoFisher) according to the manufacturer’s instructions. Cells were subjected to indicated assays after being transfected for 48 h. The DNA construct for human ACSL3 (GFP-ACSL3) was a kind gift from Albert Pol’s lab and has been described previously^73^.

#### SDS PAGE and western blot analysis

Cell lysates were determined using a BCA Protein Assay Kit as the standard. Estimated proteins (20-40 μg) were separated by SDS PAGE and transferred to PVDF membranes. Western blots were performed as standard procedures. Detection and quantification of target proteins was carried out on a scanning system (BIO-RAD, ChemiDocTM MP) using horseradish peroxidase (HRP)-conjugated secondary antibodies and ECL detection reagent.

#### Fusogenic liposome preparation

Liposomes were prepared as described previously^33^ from a lipid mixture of 1,2-dioleoyl-sn-glycero-3-phosphoethanolamine (DOPE), 1,2-dioleoyl-3-trimethylammonium-propane (chloride salt) DOTAP (both from Avanti Polar Lipids) and either 1-Stearoyl-2-Arachidonoyl-sn-glycero-3-PE (PE-AA (18:0/20:4)) or 1-Stearoyl-2-Docosahexaenoyl-sn-glycero-3-PE (PE-DHA, 18:0/22:6) (both from Cayman Chemical) at a ratio of 47.5:47.5:5. Lipid blends were in MeOH:CHCl_3_ (1:3, vol/vol). Following the generation of a thin film using a stream of nitrogen gas, the vesicles were formed by addition of 20 mM Hepes pH 7.5 (lipid concentration of 2.8 μmol/ml) and incubated for 1.5 h at room temperature. Glass beads were added to facilitate rehydration. The liposome dispersion was sonicated for 30 min (Transsonic T 310, Elma).

#### Fusogenic liposome treatment

Cells were seeded on glass coverslips in 6-well plates at 3 × 10^5^ cells/well and CAV1-mCherry expression at near endogenous levels was achieved by the addition of 0.5 ng/ml doxycycline hyclate (Sigma-Aldrich) to the media 18 h prior to liposome treatment as described previously^33^. Liposomes were added to the cells at a final concentration of 7 nmol/ml and incubated for 15 min before the cells were fixed with 3% PFA in PBS for 15 min at RT. For ACSL4 inhibition, cells were treated with 10 μM Rosiglitazone for 24 h prior to the liposome addition.

#### Quantification and Statistical Analysis

GraphPad Prism software version 10.0 was used for statistical analysis. An unpaired two-tailed Student’s *t*-test was used to determine significance between two groups. Significance in multiple groups was assessed by one-way or two-way analysis of variance (ANOVA), followed by Tukey’s or Dunnett’s test, as specified in the legends. Quantified values are presented as mean ± SD. All tests used a significance (alpha) level of 0.05. The exact P-values are reported in the figures. If the P-value is less than 0.0001, we report “****” in the figures. All the statistical details of experiments can be found in the figure legends.

## Notes

### Competing Interest Statement

The authors have declared no competing interest.

