## Supplementary figures and images for "Unsaturated lipids as key control points for caveola formation and disassembly"

### Supplemental figures

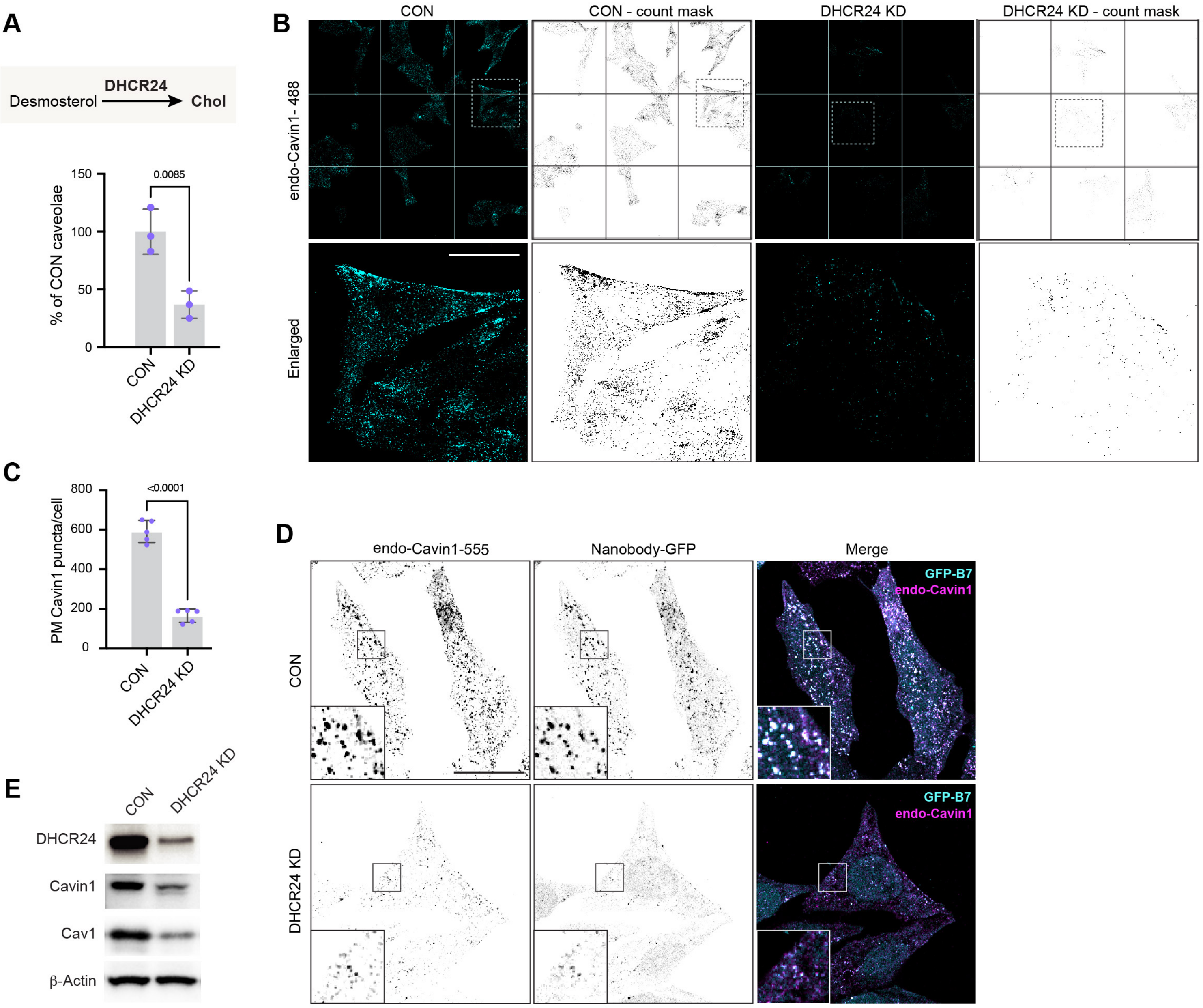

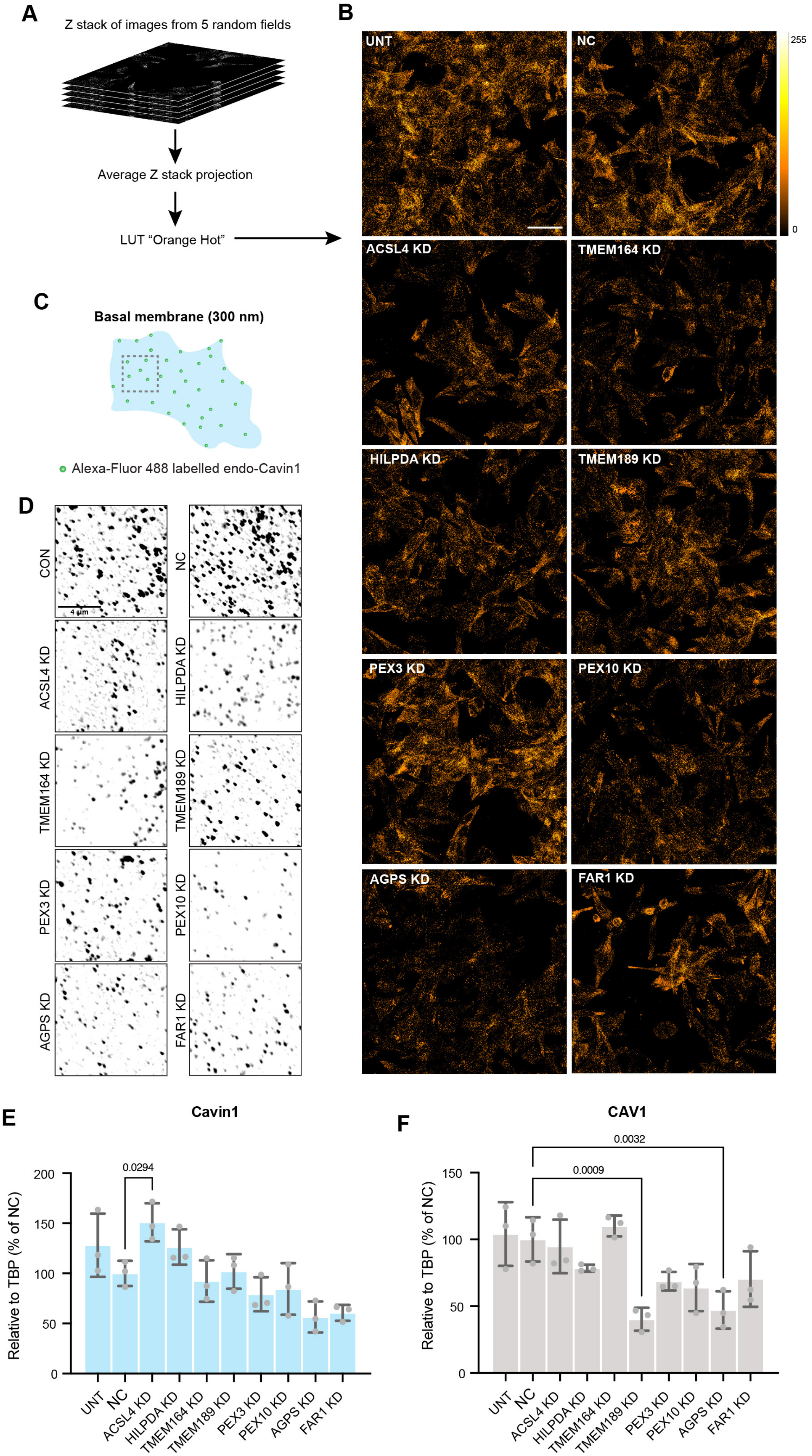

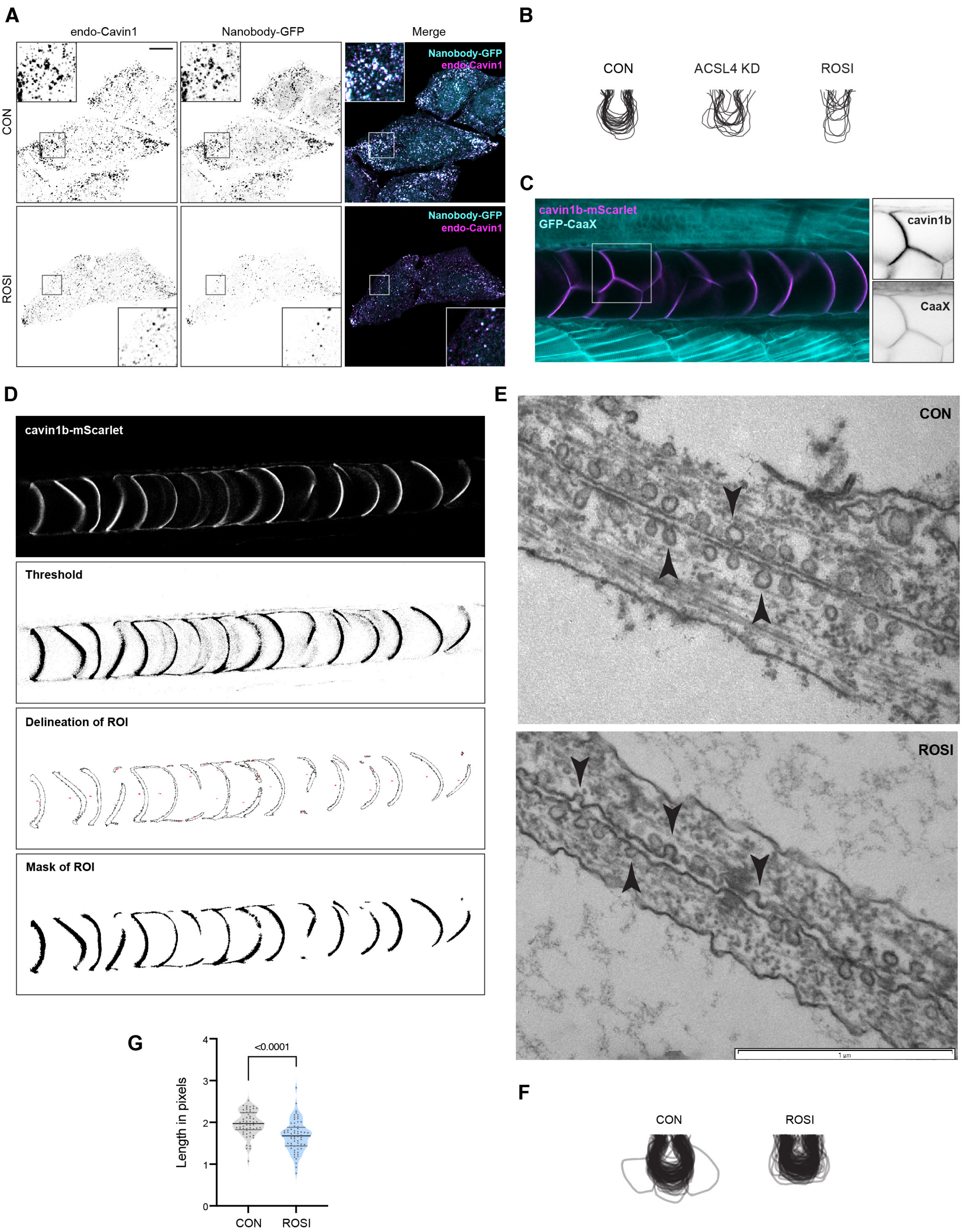

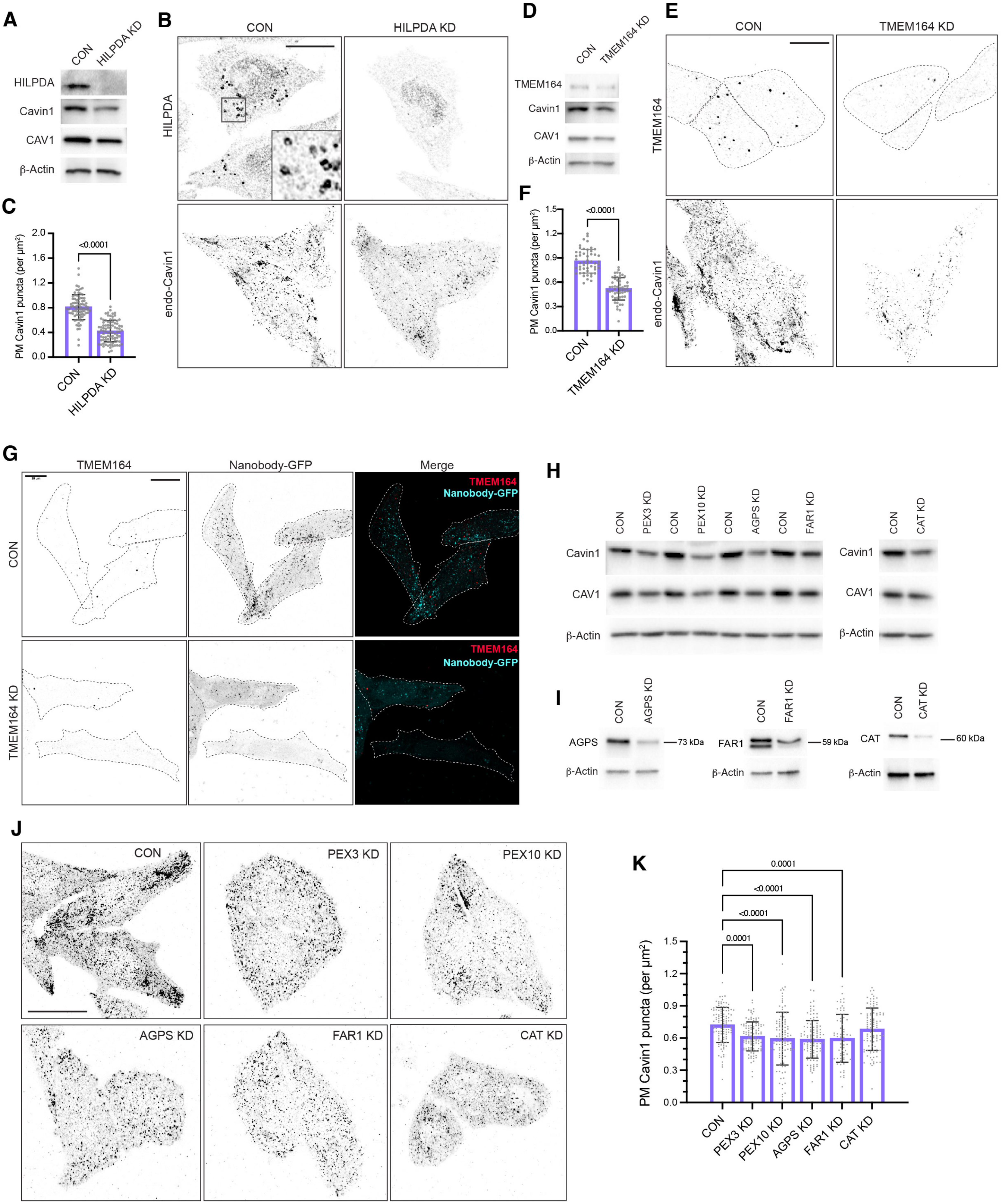

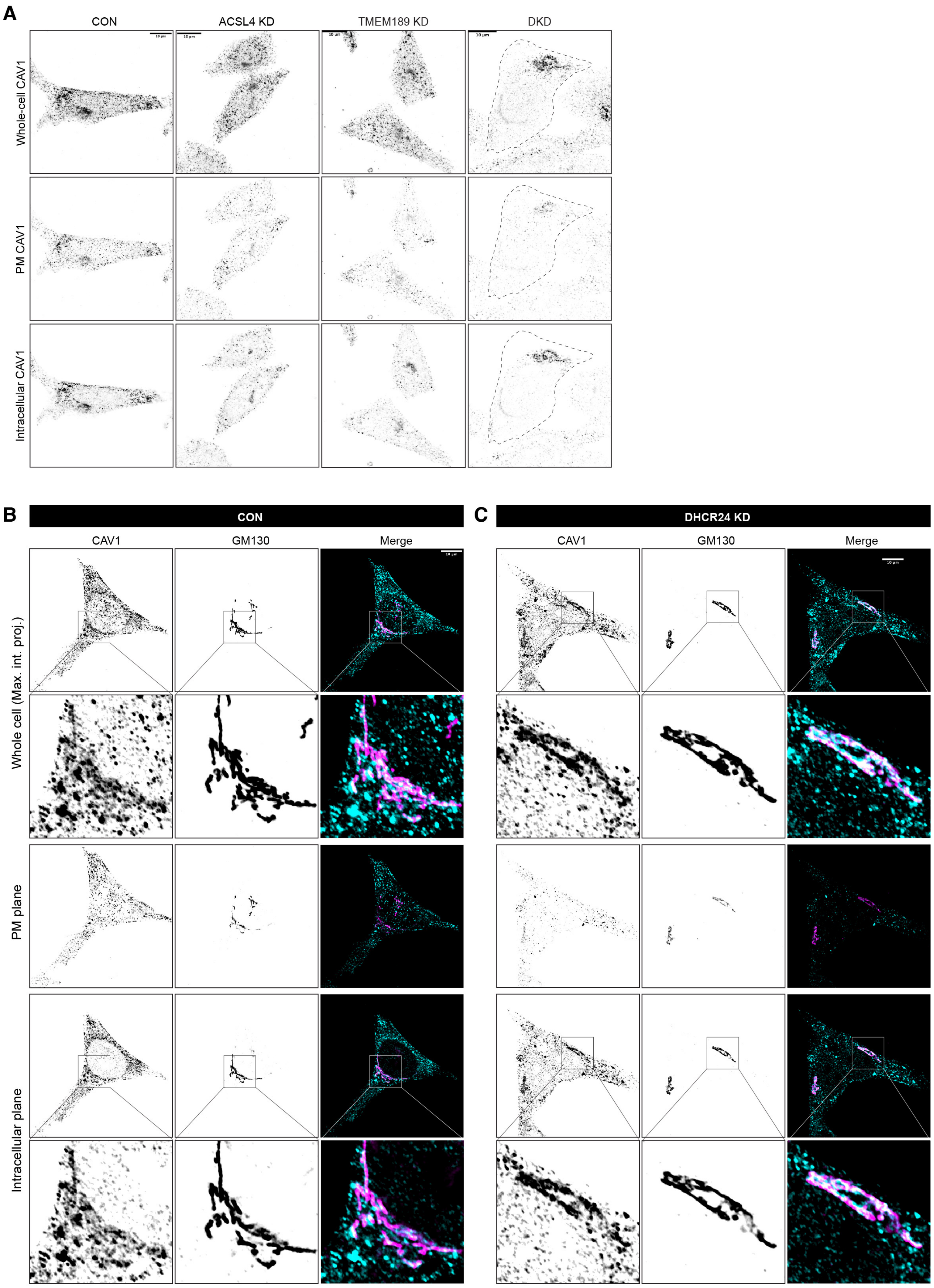

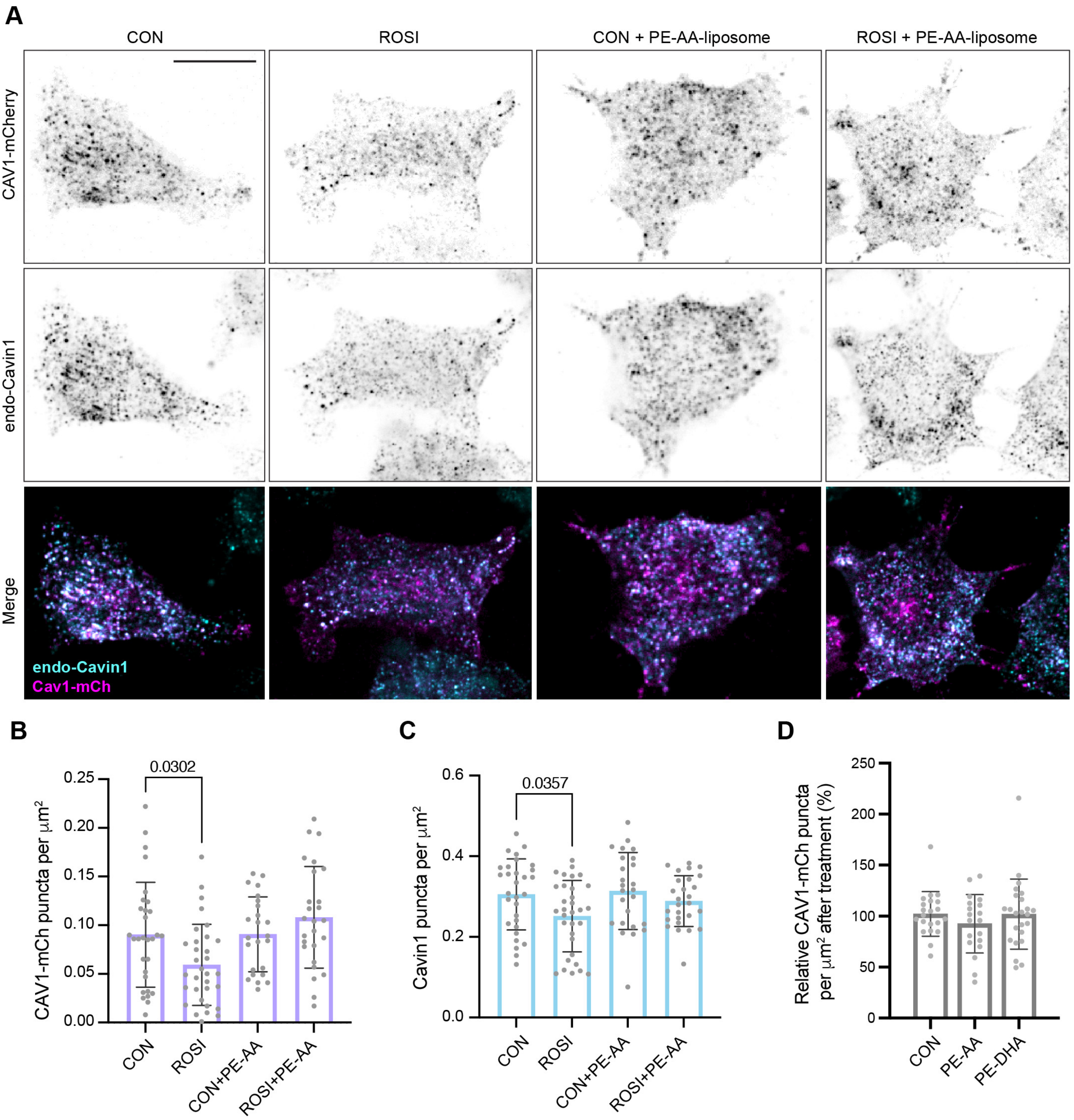
